# Membrane curvature enhances oxidation within lipid bilayers in a composition-dependent manner

**DOI:** 10.64898/2025.12.03.692219

**Authors:** Juhee Kim, Stephanie N. Bartholomew, Wade F. Zeno

## Abstract

Excessive production of reactive oxygen species (ROS) in cells results in oxidative stress, which can promote lipid oxidation in cellular membranes. This oxidation of membrane lipids accompanies various diseases and can even result in cell death through processes such as ferroptosis. The complex compositions and diverse morphologies of cellular membranes make understanding the mechanisms of lipid oxidation challenging, especially when attempting to investigate membrane composition and curvature simultaneously. Here, we utilize reconstituted lipid membranes and the fluorescent oxidation probe C11-BODIPY to quantify oxidation in lipid bilayers as functions of both lipid composition and membrane curvature. By tethering synthetic lipid vesicles to glass substrates, we were able to monitor oxidation of the fluorescent probe on a per vesicle basis using fluorescence microscopy. Our results demonstrate that highly curved membranes markedly increase both the rate and extent of oxidation across diverse membrane compositions. This effect arises from greater exposure of lipid tails to the aqueous environment, which allows more efficient transport of ROS into the hydrophobic core of the bilayer. Compositional effects on oxidation are most pronounced in membranes with low curvature (i.e., greater than 100 nm diameter) and become progressively weaker as curvature increases. We found that low to moderate cholesterol levels (i.e., 10–25 mol%) suppress curvature-dependent oxidation by tightening lipid packing, whereas high cholesterol content (i.e., 50 mol%) restores curvature sensitivity by influencing lateral lipid mobility. Together, these findings establish membrane curvature and lipid composition as interdependent determinants of oxidative susceptibility, offering new insight into how cells regulate or resist oxidative stress.

**Statement of significance:** Oxidative stress drives lipid oxidation in cellular membranes, but the combined influence of membrane curvature and composition remains poorly defined. Using reconstituted lipid vesicles and a fluorescent oxidation probe, we show that oxidation in lipid bilayers is enhanced in smaller vesicles (i.e., highly curved membranes). The oxidation rate increases with unsaturated lipids, while cholesterol suppresses this effect. Measurements of membrane packing and diffusivity support these findings, demonstrating how curvature and composition together govern membrane susceptibility to oxidative damage. These results provide new insight into the physicochemical basis of membrane stability under oxidative stress and have broad implications for understanding the vulnerability of curved cellular membranes.

## INTRODUCTION

Excessive generation of reactive oxygen species (ROS) within cells results in oxidative stress, which underlies numerous disorders such as cancer and neurodegenerative disease (1, 2). During cellular oxidative stress responses, ROS can lead to oxidation of lipids in cellular membranes, converting the hydrocarbon chains within lipid tails to aldehydes and carboxylic acids (3). These truncated lipids result in destabilization and increased permeabilization of the membrane (4–6), ultimately disrupting cellular function. In some cases, such as with ferroptosis, lipid oxidation results in cell death (7–9).

Lipid oxidation proceeds through three stages: initiation, propagation, and termination (10–12). During initiation, ROS react with unsaturated carbon double bonds in lipid tails and create lipid radicals. During propagation, lipid radicals are converted to lipid peroxyl radicals through reactions with oxygen. These reactive intermediates laterally diffuse and attack neighboring lipids, generating new radicals that spread oxidation across the membrane. The process is terminated when lipid radical species combine to form non-radical products.

Membrane composition and curvature strongly govern the susceptibility of lipids to oxidation. Polyunsaturated fatty acids (PUFAs) such as linoleic acid, one of the most abundant PUFAs in cell membranes (13), are highly susceptible to oxidation, producing reactive products that can further damage proteins (14–16). Cholesterol also modulates oxidation: it can undergo ROS- and enzyme-mediated oxidation (17, 18), which ultimately impacts neurotransmitter release (19) and promotes apoptosis (20). Beyond composition, membrane morphology is also an important consideration. Highly curved membranes, such as small unilamellar vesicles (< 100 nm diameter) undergo higher rates of oxidation than larger vesicles, likely because greater spacing between headgroups on curved membranes increase ROS accessibility (21, 22). These observations illustrate that curvature alters the physical environment of the membrane, making it essential to consider both composition and physical properties when assessing oxidative behavior. Specifically, lipid packing and lateral mobility are two physical properties that can be modulated by composition and curvature (23–25). Lipids that are unsaturated with relatively large tail groups (e.g., PUFAs) pack less densely, resulting in higher solvent accessibility to the hydrophobic membrane core (i.e., packing defects). In addition, cholesterol can affect membrane packing in a lipid saturation dependent manner, where order is increased in unsaturated membranes and decreased in saturated membranes (26). These changes in turn regulate oxidation and membrane fluidity (27). Lipid tail length and the degree of unsaturation also impact membrane fluidity, with shorter tails and greater unsaturation increasing lipid mobility (28, 29). Recent experimental work has shown that oxidation occurs more slowly when membrane fluidity is reduced (30).

Although many studies have examined how membrane composition and curvature influence lipid oxidation, these factors are usually considered in isolation. In reality, composition and morphology act together to shape physical properties that govern how membranes respond to oxidative stress. Because changes in one property often alter the other, disentangling their combined effects remains difficult. As a result, the influence of curvature and composition on lipid oxidation has not been systematically explored.

In this study, we provide a framework for understanding how curvature, composition, and the resulting membrane properties act in concert to dictate the susceptibility of membranes to oxidative damage. Here, the ROS of choice was the hydroxyl radical (OH^•^), which was generated via Fenton’s Reaction (i.e., Fe^2+^ and H_2_O_2_). To measure oxidative response to ROS in lipid membranes, we used the fluorescent oxidation probe C11-BODIPY, which permeates the membrane. This probe is known to have a sensitivity to ROS comparable to or slightly greater than that of endogenous lipids (31, 32). Using C11-BODIPY, we monitored its oxidation in reconstituted lipid vesicles with diameters ranging from approximately 40 nm to 240 nm with fluorescence confocal microscopy. Membrane fluidity and diffusivity were assessed using fluorescence anisotropy and fluorescence recovery after photobleaching (FRAP), respectively, while membrane packing was analyzed using Laurdan generalized polarization (GP). Overall, the rates and extents of oxidation exhibited notable curvature sensitivity for nearly all compositions, increasing as membranes became more curved. This effect is often presumed to arise from the higher density of packing defects (i.e. exposure of hydrophobic lipid tails) on curved membranes. To test this, we first examined dilauroylphosphatidylcholine (DLPC) vesicles, where curvature indeed enhanced oxidation as expected. We then turned to diphytanoylphosphatidylcholine (DPhPC), a lipid that forms defect-rich membranes (33), to ask whether defects alone could account for curvature-dependent oxidation. Surprisingly, despite its intrinsically high defect density, DPhPC displayed overall oxidation kinetics and extents that were nearly identical to DLPC. While DPhPC membranes had lower Laurdan GP values (i.e., greater solvent accessibility), at the same time, DPhPC membranes exhibited slower lateral diffusion, and higher anisotropy (i.e., lower molecular mobility). These results indicate that curvature-induced oxidation cannot be explained solely by packing defects but instead reflects a balance among multiple membrane properties. Moreover, we suggest that distinct membrane features can differentially influence the stages of oxidation, with ROS accessibility primarily impacting initiation and diffusional constraints shaping propagation. To further dissect this interplay, we systematically varied acyl-chain unsaturation and cholesterol content to map how each parameter modulates oxidation dynamics. Our findings demonstrate that oxidation in lipid bilayers is governed not simply by curvature-driven defects, but by the concerted interplay of lipid composition, molecular mobility, and interfacial packing, all of which together define the susceptibility of membranes to oxidative damage.

## MATERIALS AND METHODS

### Materials

HEPES (2-[4-(2-hydroxyethyl)piperazin-1-yl]ethanesulfonic acid, BP310-500), BODIPY™ 581/591 C11 (D3861) were purchased from Thermo Fisher Scientific (Waltham, MA). PLL (Poly-L-Lysine) HCl (P2658), H_2_O_2_ (Hydrogen peroxide, 30%, HX0635), FeSO_4_•7H_2_O (Iron(II) sulfate heptahydrate, F7002), and DPH (1,6-Diphenyl-1,3,5-hexatriene, D208000) were purchased from Millipore Sigma (St. Louis, MO). DLPC (1,2-dilauroyl-sn-glycero-3-phosphocholine, 850335), DPhPC (1,2-diphytanoyl-sn-glycero-3-phosphocholine, 850356), POPC (1-palmitoyl-2-oleoyl-glycero-3-phosphocholine, 850457), DOPC (1,2-dioleoyl-sn-glycero-3-phosphocholine, 850375), cholesterol (700100), DLiPC (1,2-dilinoleoyl-sn-glycero-3-phosphocholine, 850385), DSPE-PEG(2000)-Biotin (1,2-distearoyl-sn-glycero-3-phosphoethanolamine-N-[biotinyl(polyethylene glycol)-2000] (ammonium salt), 880129) were purchased from Avanti Polar Lipids (Alabaster, AL). DPPE-ATTO 647N (AD 647N-151) was purchased from ATTO-TEC (Wetzlar, Germany). mPEG-SVA (mPEG-Succinimidyl Valerate, MW 5,000, 162-130) and Biotin-PEG-SVA (MW 5,000, 164-87) were purchased from Laysan Bio, Inc (Arab, AL). Chloroform-D (D, 99.8%, DLM-7-100) was purchased from Cambridge Isotope Laboratories, Inc (Tewksbury, MA). Laurdan (CDX-D0098) was purchased from AdipoGen Life Sciences (San Diego, CA).

### Small unilamellar vesicle (SUV) preparation

SUVs were prepared by using a syringe extruder. Lipids in chloroform were mixed in a conical glass vial, dried with a N_2_ stream, and kept in a vacuum for 2 hours to remove residual solvent. To hydrate the dried lipid film, HEPES buffer (25 mM HEPES, 150 mM NaCl mM, pH 7.4) was added to the conical vial and incubated for at least 15 minutes. After thorough vortex mixing, the lipid suspensions were subjected to five freeze-thaw cycles (1 minute of freezing in liquid nitrogen and 3 minutes of thawing in a 50 °C water bath). The resulting lipid suspension was then extruded using a 100 nm pore size filter (800309, Cytiva, Wilmington, DE). To minimize oxidation of C11-BODIPY prior to experiments, sample buffers were purged with N_2_ for 10 minutes prior to lipid hydration and vials were capped under N_2_. The size of the extruded vesicles was measured by dynamic light scattering (DLS), which is summarized in Table S1.

### Confocal microscopy imaging of SUVs

SUVs were imaged using a tethered vesicle assay on a STELLARIS 5 laser scanning confocal microscope (Leica Microsystems). The detailed experimental protocol has been described previously (30). Briefly, a PLL-PEG-Biotin complex was passivated onto a Hellmanex-cleaned coverslip and incubated with neutravidin. SUVs containing phospholipids, C11-BODIPY, DPPE-ATTO 647N, and DSPE-PEG(2000)-Biotin were tethered within imaging wells and subsequently exposed to 7.5 μM H_2_O_2_ and 0.15 μM FeSO_4_ to induce oxidation via hydroxyl radicals. Imaging wells subjected to oxidizing conditions were incubated with hydroxyl radicals for either 5, 10, or 30 minutes. Excitation lasers at 638 nm and 488 nm were used to excite ATTO 647N and C11-BODIPY, respectively. The detection wavelength ranges were 645–749 nm for ATTO 647N, 570–620 nm for unoxidized C11-BODIPY, and 500–540 nm for oxidized C11-BODIPY. The fluorescence intensities of individual SUVs were analyzed using CME analysis (34).

### Fluorescence recovery after photobleaching

For FRAP analysis, planar supported lipid bilayers were used. Both unilamellar bilayers (single bilayer or SBL) and multilamellar bilayers (multi-bilayers or MBLs) were used. To create SBLs, SUVs composed of PC lipid (99.875 mol%) and DPPE-ATTO 647N (0.125 mol%) were coated onto glass cover slips. For samples containing saturated lipids (i.e., DLPC or DPhPC), SUVs were synthesized by drying chloroform-dissolved lipid mixtures with a N_2_ stream then placing them under vacuum for 2 hours to remove the residual solvent. The lipid films were then hydrated using HEPES buffer (25 mM HEPES, 150 mM NaCl, pH 7.4) that was purged with N_2_ for 10 minutes, then sonicated using a probe tip sonicator. A total of four sonication cycles (2 minutes of sonication and 2 minutes of rest) were performed for each sample on ice. Sonicated lipids samples were then centrifuged at 21,100 Xg for 10 minutes to remove tip shards. For samples containing unsaturated lipids (i.e., DLiPC, DOPC, or POPC), SUVs were prepared using the extrusion methods described above (under “SUV preparation”). For SBL formation, SUVs were diluted to the final concentration of 100 μM total lipid, incubated in an imaging well on top of a Hellmanex-cleaned glass coverslip for 10 minutes, and rinsed five times with HEPES buffer (25 mM HEPES, 150 mM NaCl, pH 7.4) to remove excess lipids.

To create MBLs, lipid mixtures consisting of PC lipid (99.875 mol%) and DPPE-ATTO 647N (0.125 mol%) in chloroform were prepared in conical vials then dried down with a N_2_ stream to remove solvent. Afterward, a hexane:methanol (97:3, volume ratio) solution was added to the dried lipid film to resolubilize the lipids. Depending on the lipid mixture used, the final concentration of total lipids ranged from 6.6 mM to 10.9 mM. The solubilized lipid solution was then spin coated onto a Hellmanex-cleaned round coverslip (No. 1.5, diameter of 25 mm) for 40 seconds with 3,000 rpm. The coated coverslip was kept under vacuum for 2 hours to remove the remaining solvent and assembled into a Sykes-Moore Chamber (1943-11111, Bellco glass, Vineland, NJ). To hydrate the lipid film, 500 uL of HEPES buffer (25 mM HEPES, 150 mM NaCl, pH 7.4) was added to the chamber prior to imaging.

Prior to photobleaching the region of interest, 3 image frames were taken, and FRAP was conducted using a 638 nm laser at 100% intensity over a circular region with 15 μm diameter. Afterward, a minimum of 150 image frames (0.648 second intervals) were acquired to monitor fluorescence recovery. The detection wavelength range used for ATTO 647N was 645–749 nm and intensity distribution for each frame was quantified using ImageJ. Diffusion coefficients were calculated using a uniform circular beam model (35, 36). A detailed explanation of this method is provided in the Supporting Discussion.

### Fluorescence spectroscopy

SUVs containing 0.05 mol% of C11-BODIPY were prepared as described above in the “SUV preparation” section. SUVs (200 μM total lipid concentration) were oxidized using 20 mM H_2_O_2_ and 1 mM FeSO_4_ in buffer consisting of 25 mM HEPES, 150 mM NaCl, and 1 mM EGTA (pH 7.4). Fluorescence emission spectra of C11-BODIPY were obtained every 10 seconds at 23 °C over the 500–650 nm range using 488 nm excitation on a spectrofluorometer (JASCO FP-8500).

### Fluorescence anisotropy

Stock solutions of DPH in ethanol:DMSO (97:3 volume ratio, 200 µM) were added to extruded SUV solutions (100% PC lipid, 60 µM total lipid) such that the molar ratios of lipid:DPH were 100:1. DPH was excited at 355 nm and fluorescence emission was monitored at 425 nm using a spectrofluorometer (JASCO FP-8500).

### Laurdan generalized polarization

SUVs containing 0.2 mol% Laurdan (100 μM total lipid concentration) were prepared using 200 nm filters (10417004, Cytiva, Wilmington, DE) as described above (under “SUV preparation”). Laurdan was excited at 340 nm and emission spectra were obtained from 400–550 nm at 23 °C using a spectrofluorometer (JASCO FP-8500). The fluorescence emission spectra were normalized to their maximum fluorescence intensities, and the GP values were calculated using equation 1, where I_440_ and I_490_ refer to the normalized fluorescence intensity at 440 nm and 490 nm, respectively.

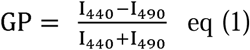

### ^1^H Nuclear magnetic resonance (NMR) spectroscopy

SUVs composed of 100% PC lipids were prepared using 100 nm extrusion filters as described above (under “SUV preparation”). Lipid concentrations for NMR spectroscopy samples were chosen such that they were consistent with their experimental counterparts in the tethered vesicle assays. Detailed calculations for determining these lipid concentrations are provided in the Supporting Discussion. Solutions containing SUVs were incubated with the appropriate concentrations of Fenton’s Reagents (i.e., FeSO_4_ and H_2_O_2_) for a minimum of 10 minutes while stirring. Afterward, the reaction mixtures were combined with hexane:IPA (3:1 volume ratio) and centrifuged for 10 minutes at 3,400 Xg to extract lipids into the hexane layer. After centrifugation, the top hexane layer was extracted into a separate vial, dried down with a N_2_ stream, and resolubilized in CDCl_3_. Using a standard of CHCl_3_ (δ 7.26) and 400 MHz field strength, ^1^H NMR spectra were obtained using an NMR spectrometer (Varian Mercury 400).

## RESULTS

### Impact of membrane curvature on oxidation

Using the fluorescent oxidation probe C11-BODIPY, we first monitored oxidation of the probe in membranes composed primarily of DLPC. C11-BODIPY contains a conjugated diene group that serves as an oxidation site (Figure 1A, left) (31). Upon oxidation by hydroxyl radicals, the phenyl group is cleaved, creating a carboxyl moiety that shortens the conjugation system (Figure 1A, right) (37). As a result, the maximum peak of the C11-BODIPY emission spectrum shifts from 590 nm to 520 nm, enabling ratiometric quantification of its oxidized and unoxidized states (Figure 1B). C11-BODIPY is known to be predominantly located in the shallow region of the membrane (31, 38). Upon oxidation, the probe increases in polarity, causing it to move slightly towards the bilayer headgroups at the water interface (31). Fenton’s Reagent (i.e., a mixture of hydrogen peroxide and FeSO_4_) was used to generate hydroxyl radicals and induce oxidation of the probe in the membrane. To monitor the oxidation of C11-BODIPY on a per vesicle basis, we utilized a tethered vesicle assay (Figure 1C). Confocal micrographs reveal clear colocalization among all three fluorescence channels of interest: the tracer dye ATTO 647N, the 590 nm emission for C11-BODIPY in the unoxidized state, and the 520 nm emission for C11-BODIPY in the oxidized state (Figure 1D). The fluorescence intensity of ATTO 647N was used to determine vesicle diameters via quantitative calibrations with DLS measurements (Figure S1). The fluorescence emission intensities at 590 nm and 520 nm were used to determine the relative extent of oxidation in each SUV. When looking at the merged channel, upon 30 minutes of exposure to hydroxyl radicals, the vesicles became greener in the merged fluorescence channel (Figure 1D, Merged), visually indicating an increase in the relative fluorescence emission at 520 nm relative to 590 nm. This result corresponds to an increase in the proportion of C11-BODIPY in the oxidized state versus the unoxidized state. Next, we used emission spectra from C11-BODIPY samples with known fractions of oxidized and unoxidized populations to generate quantitative calibrations (Supporting Discussion, Figure S2, Table S2). These calibrations allowed us to convert the emission intensities from each tethered vesicle into the corresponding molar fraction of C11-BODIPY in the oxidized state (Figure 1E). Here, vesicle populations were sorted and binned by diameter. Importantly, the 175 nm population is not absent; vesicles in this size range are included within the 200 nm bin, which spans diameters from 160–240 nm. Interestingly, prior to the addition of ROS, the extent of oxidation was not uniform across vesicle sizes and increased monotonically with decreasing vesicle diameter. This background oxidation likely arises during vesicle preparation, where extrusion and freeze–thaw cycles occur in the presence of dissolved oxygen. The higher fraction oxidized in smaller vesicles suggests that highly curved membranes are intrinsically more susceptible to oxidation, potentially due to enhanced lipid splay. This background oxidation was minimized by preparing vesicles with N_2_-purged buffers (see methods). For each diameter, the fraction of oxidized C11-BODIPY increased over time (Figures 1E-F). Additionally, the molar fraction of oxidized C11-BODIPY increased monotonically with vesicle curvature at all time points (Figure S3). To calculate the initial rate of oxidation, the fractions of oxidized C11-BODIPY at first three time points (0, 5, and 10 minutes) were linearly regressed (Figure 1G). This initial rate of oxidation also exhibited a strong curvature preference, with the fastest rate occurring on the smallest vesicles (Figure 1H).

**Figure 1.**
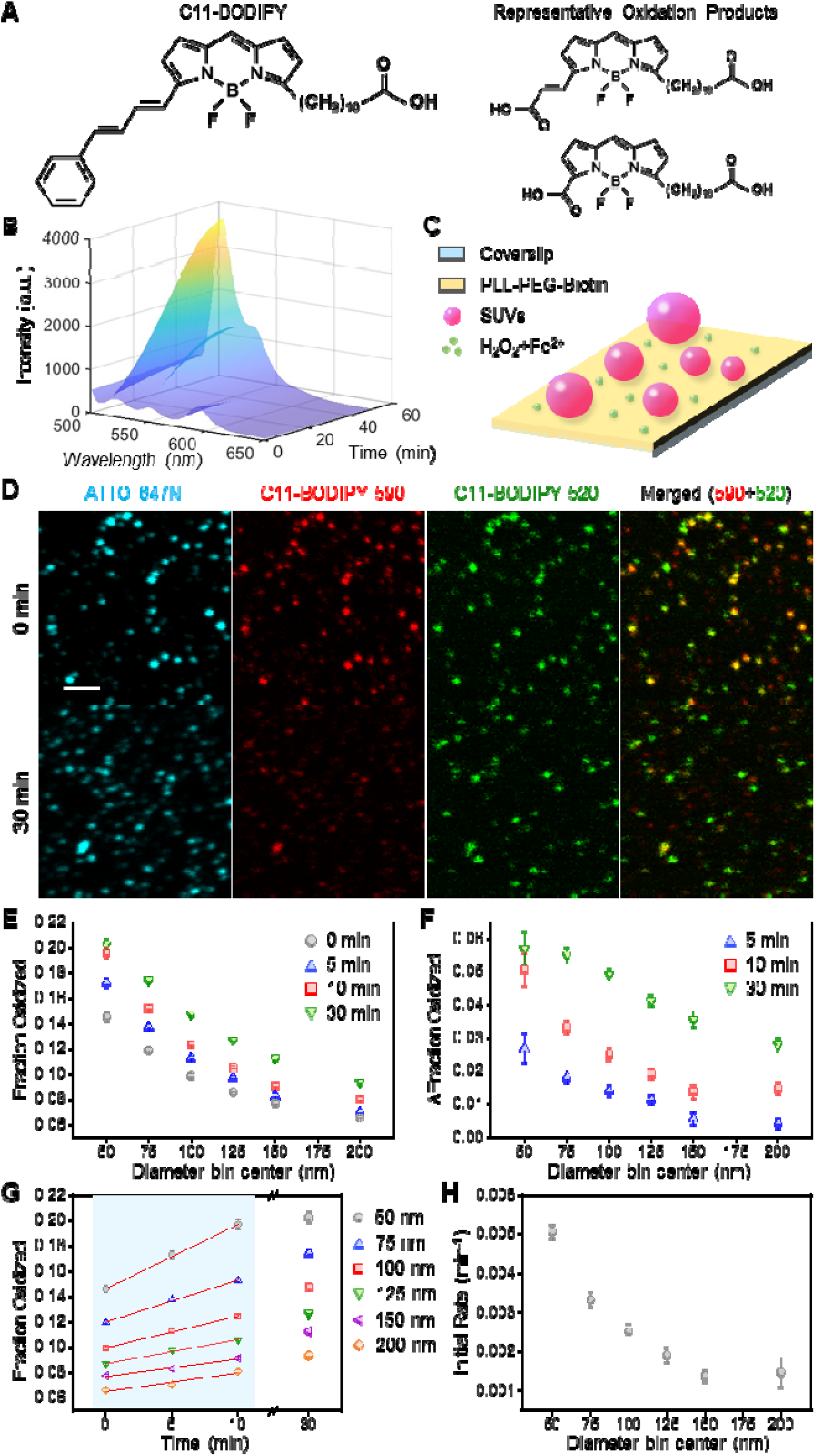
Membrane curvature enhances oxidation in lipid bilayers. **A)** Structure of C11-BODIPY (left) and representative oxidation products (right). **B)** Emission spectrum of C11-BODIPY during the course of oxidation. **C)** Schematic of the tethered vesicle assay. **D)** Representative fluorescence micrographs of SUVs tethered on a cover slip and exposed to oxidizing conditions for 30 minutes (scale bar: 2 μm). **E)** The average molar fraction of oxidized C11-BODIPY in each diameter bin for different experimental time points. **F)** The change in the molar fraction of oxidized C11-BODIPY relative to its initial value for each diameter bin. **G)** The average molar fraction of oxidized C11-BODIPY over time for each diameter bin. The shaded region represents the values over which linear regressions (red lines) were performed when calculating initial rates of oxidation. **H)** The initial rate of oxidation as a function of vesicle diameter. Vesicles used in panel **B** were composed of 99.95 mol% DLPC and 0.05 mol% C11-BODIPY (200 µM total lipid concentration), and oxidation was induced using 20 mM H_2_O_2_ and 1 mM FeSO_4_. All vesicles used in panels **D-H** were composed of DLPC (98.5 mol%), C11-BODIPY (0.5 mol%), DPPE-ATTO 647N (0.5 mol%), and DSPE-PEG(2000)-Biotin (0.5 mol%), and oxidation was induced using 7.5 μM H_2_O_2_ and 0.15 μM FeSO_4_. The number of SUVs in each diameter bin in **E**-**H** varied from 906–4641. Data points in **E-H** represent the average value of the bin, and the error bars correspond to the standard error of the mean for each bin. Each value displayed for a bin (i.e., 50 nm, 75 nm, 100 nm, etc.) represents the midpoint of the bin. All bins were symmetric, with endpoints aligned to the start of the next bin, ensuring no overlap.

### Impact of lipid packing defects on oxidation

The observed curvature dependence of oxidation in Figure 1 could be attributed to the increased lipid splay in highly curved vesicles, which would increase hydrophobic tail exposure to the solution environment (i.e., lipid packing defects). This increased exposure would allow a greater flux of ROS into the membrane environment. Therefore, we next investigated DPhPC membranes, which have been shown to contain higher hydrophobic tail exposure than DLPC membranes due to the presence of four branched methyl groups in the lipid tail (Figure 2A) (33). To compare the membrane packing of the two lipids, we used Laurdan, a membrane-permeable fluorescent probe that is sensitive to the polarity of its surrounding environment (Figures 2B-C) (39). In loosely packed membranes, increased water penetration and mobility at the lipid interface enhances dipolar relaxation of the water molecules near Laurdan. This relaxation causes a red shift in the Laurdan emission spectrum, moving the maximum intensity peak from 440 nm to 490 nm (40). Emission peaks closer to 440 nm yield higher GP values while peaks closer to 490 nm yield lower values (Equation 1). Therefore, membrane packing can be quantified by calculating GP values from emission spectra, where higher GP values indicate a more packed membrane. Consistent with previous studies (41, 42), the GP value of DLPC was higher than that of DPhPC, corroborating the greater hydrophobic tail exposure of DPhPC.

**Figure 2.**
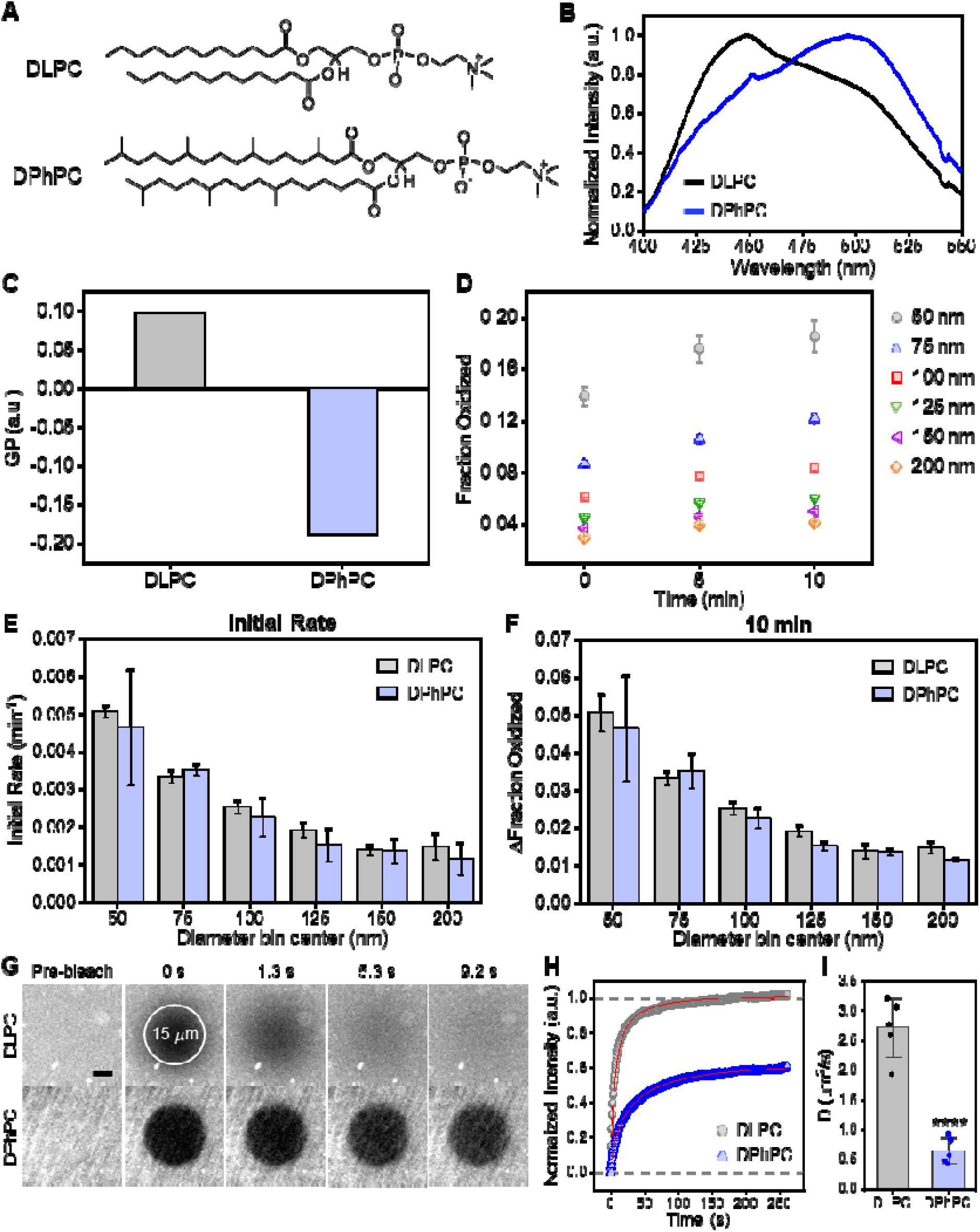
Interplay between packing defects and diffusivity governs oxidation in DLPC and DPhPC membranes. **A)** Structure of DLPC and DPhPC. **B)** The fluorescence emission spectra of Laurdan for each lipid composition. **C)** The GP value of Laurdan for each lipid composition. **D)** The average fraction of oxidized C11-BODIPY over time for each diameter bin. **E)** The initial rate of oxidation as a function of vesicle diameter. **F)** The change in the fraction of oxidized C11-BODIPY after 10 minutes of oxidation, relative to its initial value for each diameter bin. **G)** Representative fluorescence micrographs of MBLs before and after photobleaching (scale bar: 5 μm). **H)** Fluorescence recovery in MBLs over time. The red lines represent the lines of best fit from equation S7. **I)** Diffusion coefficients for each lipid composition. All vesicles used in panel **B** and **C** were composed of DLPC or DPhPC (99.8 mol%) and Laurdan (0.2 mol%). All vesicles used in panel **D-F** were composed of DPhPC (98.5 mol%), C11-BODIPY (0.5 mol%), DPPE-ATTO 647N (0.5 mol%), and DSPE-PEG(2000)-Biotin (0.5 mol%), and oxidation was induced using 7.5 μM H_2_O_2_ and 0.15 μM FeSO_4_. The number of DPhPC SUVs in each diameter bin varied from 140–4959. Data points or bars in **D-F** represent the average value of the bin and the error bars correspond to the standard error of the mean for each bin. Each value displayed for a bin represents the midpoint of the bin. All bins were symmetric, with endpoints aligned to the start of the next bin, ensuring no overlap. MBLs used in panel **G-I** were composed of DLPC or DPhPC (99.875 mol%) and DPPE-ATTO 647N (0.125 mol%). Data points in **H** represent the average normalized fluorescence intensity. Bars in **I** represent the mean and the error bars correspond to the standard deviation (*N*=5, unpaired, two-sample t-test, ****P<0.0001).

The tethered vesicle assay was used to monitor oxidation in SUVs composed primarily of DPhPC (Figure 2D). Similar to DLPC vesicles used in Figure 1G, DPhPC vesicles exhibited monotonic responses to membrane curvature at all experimental time points, with the level of oxidation increasing with increased membrane curvature. However, much to our surprise, the levels and rates of C11-BODIPY oxidation in DPhPC vesicles were not substantially different from those of DLPC vesicles (Figures 2E-F), despite DPhPC membranes having much higher exposure of hydrophobic tails. The initial rates of C11-BODIPY oxidation in DPhPC membranes were similar to the rates in DLPC membranes for all vesicle diameter bins (Figure 2E). These trends were also reflected in the quantified molar fraction of oxidized C11-BODIPY after 10 minutes of exposure to hydroxyl radicals (Figure 2F).

We conducted FRAP on MBLs composed of DLPC or DPhPC to probe differences in the lateral diffusivity of the two membranes. A fast fluorescence recovery was observed in DLPC membranes, whereas DPhPC membranes exhibited significantly slower fluorescence recovery (Figures 2G-H). This trend of a slower recovery for DPhPC compared to DLPC is quantitatively consistent with previous measurements (43, 44). The diffusion coefficient of DPhPC was approximately 4 times smaller than that of DLPC (Figure 2I). Qualitatively similar diffusion results were obtained using lipid SBLs (Figure S4) and further corroborated via fluorescence anisotropy measurements (Figure S5). Therefore, the enhancing effect of lipid packing defects on oxidation may have been offset by the inhibitory effect of reduced lipid mobility, rationalizing the similar oxidation behavior observed for C11-BODIPY in DLPC and DPhPC vesicles.

### Impact of lipid monounsaturation on oxidation

So far, we measured the oxidation of C11-BODIPY in membranes made of saturated lipids. To understand the effect of unsaturated lipids on oxidation, we first incorporated lipids with monounsaturated oleic acid tails: palmitoyloleoylphosphatidylcholine (POPC) and dioleoylphosphatidylcholine (DOPC) (Figure 3A). In POPC membranes, the emission intensity of Laurdan at 490 nm was higher than that in DLPC membranes (Figure S6A). Accordingly, the GP value of Laurdan was lower in POPC membranes than in DLPC membranes (Figure 3B), implying reduced lipid packing in POPC due to the presence of a single unsaturated bond. The GP value decreased further in DOPC membranes, indicating an additional reduction in lipid packing associated with the second unsaturated bond. This trend is consistent with previous measurements comparing GP values in POPC and DOPC membranes (45–47). DPhPC membranes maintained the lowest GP value (Figure 3B), placing POPC and DOPC at intermediate packing levels between DLPC and DPhPC. Using FRAP on lipid MBLs (Figures 3C, S6B-C), the lateral diffusion coefficients were found to be higher in DOPC membranes than in POPC membranes, consistent with previous results (48, 49). For both DOPC and POPC membranes, the diffusion coefficients were higher than those in DPhPC membranes but lower than those in DLPC membranes. Similar results were observed using FRAP on lipid SBLs (Figure S4), as well as fluorescence anisotropy in SUVs (Figure S5) where membrane fluidity followed the trend: DLPC > DOPC > POPC > DPhPC.

**Figure 3.**
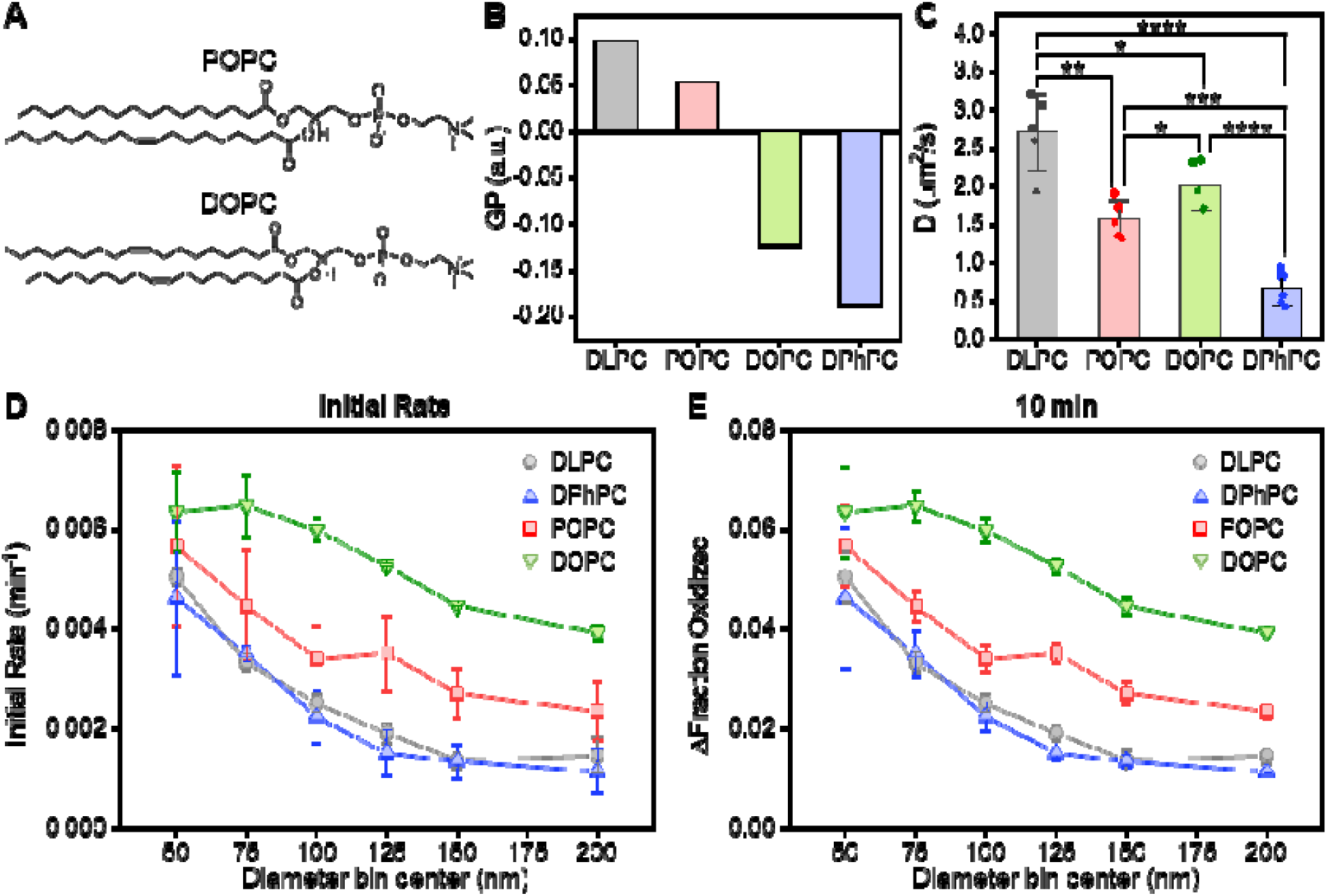
Monounsaturated lipids enhance oxidation in lipid bilayers. **A)** Structure of POPC and DOPC. **B)** The GP value of Laurdan within each lipid bilayer. **C)** Lateral diffusion coefficients for each membrane composition. **D)** The initial rate of oxidation as a function of vesicle diameter. **E)** The change in the fraction of oxidized C11-BODIPY after 10 minutes of oxidation, relative to its initial value for each diameter bin. Vesicles used in panel **B** were composed of PC lipids (99.8 mol%) and Laurdan (0.2 mol%). MBLs used in panel **C** were composed of PC lipids (99.875 mol%) and DPPE-ATTO 647N (0.125 mol%). Bars in **C** represent the mean and the error bars correspond to the standard deviation (*N*=5, unpaired, two-sample t-test, *P<0.05, **P<0.01, ***P<0.001, ****P<0.0001). All vesicles used in panel **D** and **E** were composed of PC lipids (98.5 mol%), C11-BODIPY (0.5 mol%), DPPE-ATTO 647N (0.5 mol%), and DSPE-PEG(2000)-Biotin (0.5 mol%), and oxidation was induced using 7.5 μM H_2_O_2_ and 0.15 μM FeSO_4_. The number of SUVs in each diameter bin varied from 680–6644 (POPC), and 462–8402 (DOPC). Data points in **D** and **E** represent the average value of the bin, and the error bars correspond to the standard error. Each value displayed for a bin represents the midpoint of the bin. All bins were symmetric, with endpoints aligned to the start of the next bin, ensuring no overlap.

We used Fenton’s reagent concentrations that were sufficient to oxidize C11-BODIPY directly but not high enough to oxidize double bonds in POPC or DOPC. This experimental design restricted initial oxidation to C11-BODIPY alone, ensuring that POPC and DOPC could only become oxidized through propagation reactions within the membrane. ^1^H NMR spectroscopy confirmed that the chemical structures of POPC and DOPC remained unchanged after at least 10 minutes of exposure to the oxidizing conditions used (Figures S7, S8). Upon oxidation of C11-BODIPY in SUVs, we found that oxidation rates were highest in DOPC membranes for all membrane curvatures (Figures 3D, S9A-B). Oxidation rates in POPC membranes were higher than those in DLPC and DPhPC membranes. Similar trends were observed when analyzing the fraction of oxidized C11-BODIPY in SUVs after 5 minutes (Figure S9C) or 10 minutes (Figure 3E) of ROS exposure. Interestingly, the disparities in oxidation levels were minimized when curvature was greatest (Figures 3D-E, S9C).

### Impact of lipid polyunsaturation on oxidation

Polyunsaturated lipids are more susceptible to oxidation than monounsaturated lipids (10), therefore we investigated the simultaneous impacts of lipid unsaturation and membrane curvature on oxidation in lipid bilayers. Here the PUFA-containing lipid of choice was dilinoleoylphosphatidylcholine (DLiPC) (Figure 4A). Three membrane compositions were compared: membranes composed primarily of DOPC, membranes composed of DLiPC and DPhPC at a 1:1 molar ratio (50% DLiPC), and membranes composed of 3:1 DLiPC:DPhPC (75% DLiPC). With this experimental design, the average number of double bonds was held constant between DOPC and 50% DLiPC membranes at 2 double bonds per phospholipid but was increased to 3 double bonds per phospholipid in 75% DLiPC membranes. The GP values for membranes containing 50% and 75% DLiPC were similar to one another, but lower than that of DOPC (Figures 4B, S10A), likely due to the propensity of PUFA-containing bilayers to pack loosely. This reduced packing is consistent with previous reports showing that DLiPC membranes exhibit GP values comparable to or lower than DOPC membranes (46, 50, 51). The inclusion of branched-tail DPhPC lipids in the 50% and 75% DLiPC mixtures may have further decreased the GP values relative to DOPC. FRAP analysis revealed that lipid lateral diffusivity was lowest in 50% DLiPC membranes (Figures 4C, S10B). Because acyl tail unsaturation is generally associated with increased lateral diffusivity (52), this reduction indicates that the presence of DPhPC is responsible for the decreased diffusivity (Figure 2I).

**Figure 4.**
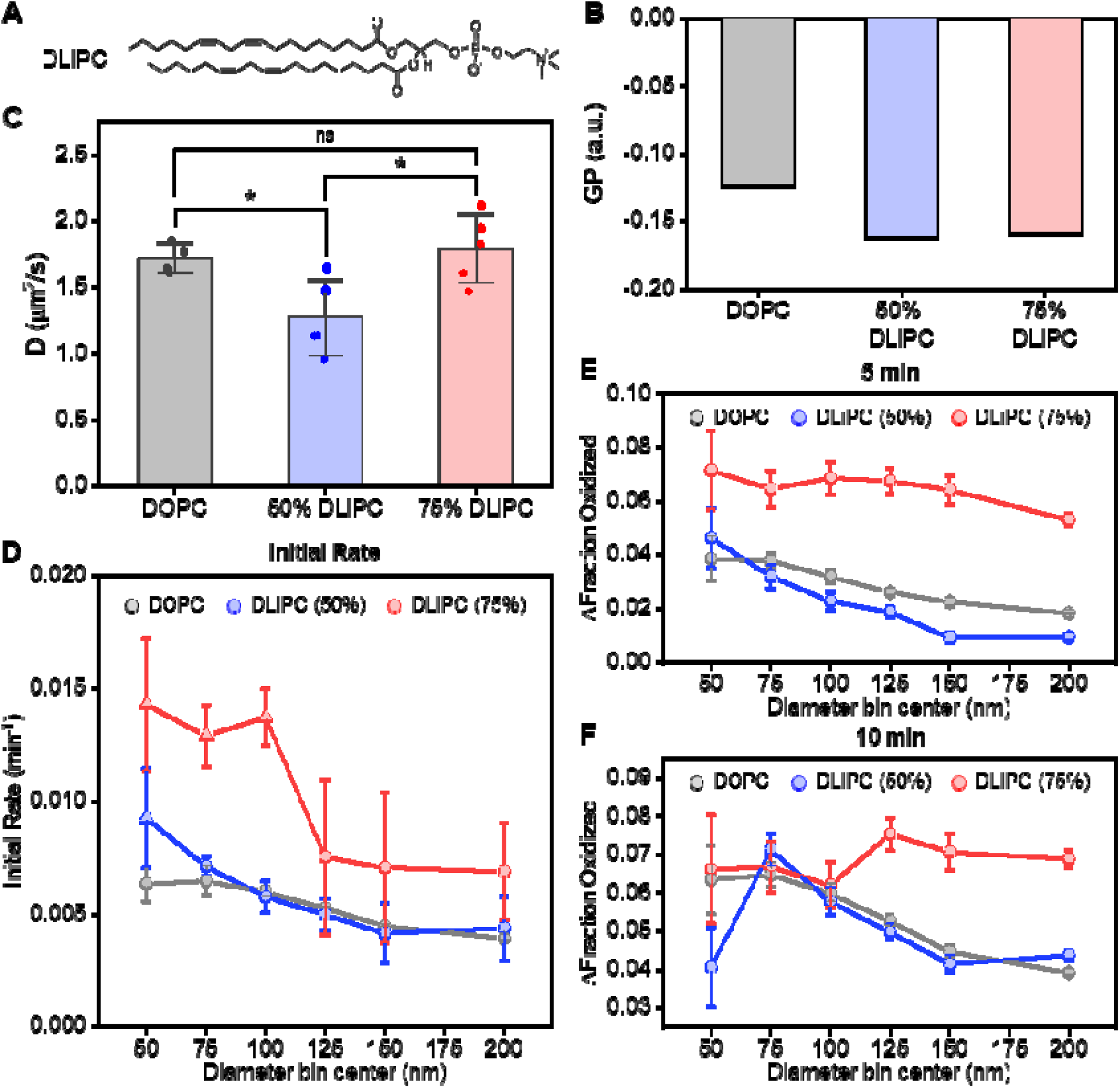
Lipid polyunsaturation further enhances oxidation in lipid bilayers. **A)** Structure of DLiPC. **B)** The GP value of Laurdan within each lipid bilayer. **C)** Lateral diffusion coefficients for each membrane composition. **D)** The initial rate of oxidation as a function of vesicle diameter. The change in the fraction of oxidized C11-BODIPY after **E)** 5 minutes and **F)** 10 minutes of oxidation, relative to its initial value for each diameter bin. Vesicles used in panel **B** were primarily composed of DOPC, 1:1 DLiPC:DPhPC (50% DLiPC), or 3:1 DLiPC:DPhPC (75% DLiPC), and Laurdan (0.2 mol%). SBLs used in panel **C** were primarily composed of DOPC, 1:1 DLiPC:DPhPC (50% DLiPC), or 3:1 DLiPC:DPhPC (75% DLiPC), and contained 0.125 mol% DPPE-ATTO 647N. Bars in **C** represent the mean and the error bars correspond to the standard deviation (*N*=4 for DOPC, and *N*=5 for 50% DLiPC and 75% DLiPC, unpaired, two-sample t-test, ns: non-significance, *P<0.05). All vesicles used in panel **D-F** were primarily composed of DOPC, 1:1 DLiPC:DPhPC (50% DLiPC), or 3:1 DLiPC:DPhPC (75% DLiPC), and contained C11-BODIPY (0.5 mol%), DPPE-ATTO 647N (0.5 mol%), and DSPE-PEG(2000)-Biotin (0.5 mol%). Oxidation was induced using 7.5 μM H_2_O_2_ and 0.15 μM FeSO_4_. The number of SUVs in each diameter bin varied from 320–4502 (50% DLiPC), and 263–3100 (75% DLiPC). Data points in **D** represent the average initial rate of oxidation calculated from linear regression of three time points (0, 5, and 10 minutes, circle) or slope of the first two time points (0 and 5 minutes, triangle) and the error bars correspond to the standard error calculated from linear regression (circle) or propagated error from the standard error (triangle). Data points in **E** and **F** represent the average value of the bin, and the error bars correspond to the standard error of difference. Each value displayed for a bin represents the midpoint of the bin. All bins were symmetric, with endpoints aligned to the start of the next bin, ensuring no overlap.

All membrane compositions elicited increased oxidation of C11-BODIPY with increased curvature, as expected (Figures 4D, S10C-D). The initial rate of oxidation was highest in 75% DLiPC membranes, which contained the highest density of double bonds (Figure 4D). The initial oxidation rate in 50% DLiPC membranes was comparable to that in DOPC membranes. These trends were qualitatively reflected when analyzing the change in fraction of oxidized C11-BODIPY after 5 minutes of ROS exposure (Figure 4E). After 10 minutes of oxidation, the change in the molar fraction of oxidized C11-BODIPY was comparable for all compositions on the smaller vesicle populations (i.e., < 100 nm diameter), but highest for 75% DLiPC compositions on the larger vesicle populations (Figure 4F).

### Impact of cholesterol content on oxidation

Cholesterol is known to order lipid tails and decrease the permeability of membranes composed of unsaturated lipids (26, 40), which in turn, decreases the interfacial exposure of hydrophobic lipid tails. To probe the impact of cholesterol on oxidation in lipid bilayers, DOPC membranes containing varying cholesterol concentrations were prepared. Laurdan fluorescence confirmed that this was the case in our experiments as GP values increased monotonically with increasing cholesterol mole fractions (Figures 5A, S11A), consistent with previous reports of Laurdan polarization in cholesterol-containing membranes (53). Also consistent with the tighter lipid packing, lateral diffusivity decreased with increasing cholesterol mole percentage (Figures 5B, S11B), in agreement with previously reported measurements (54).

**Figure 5.**
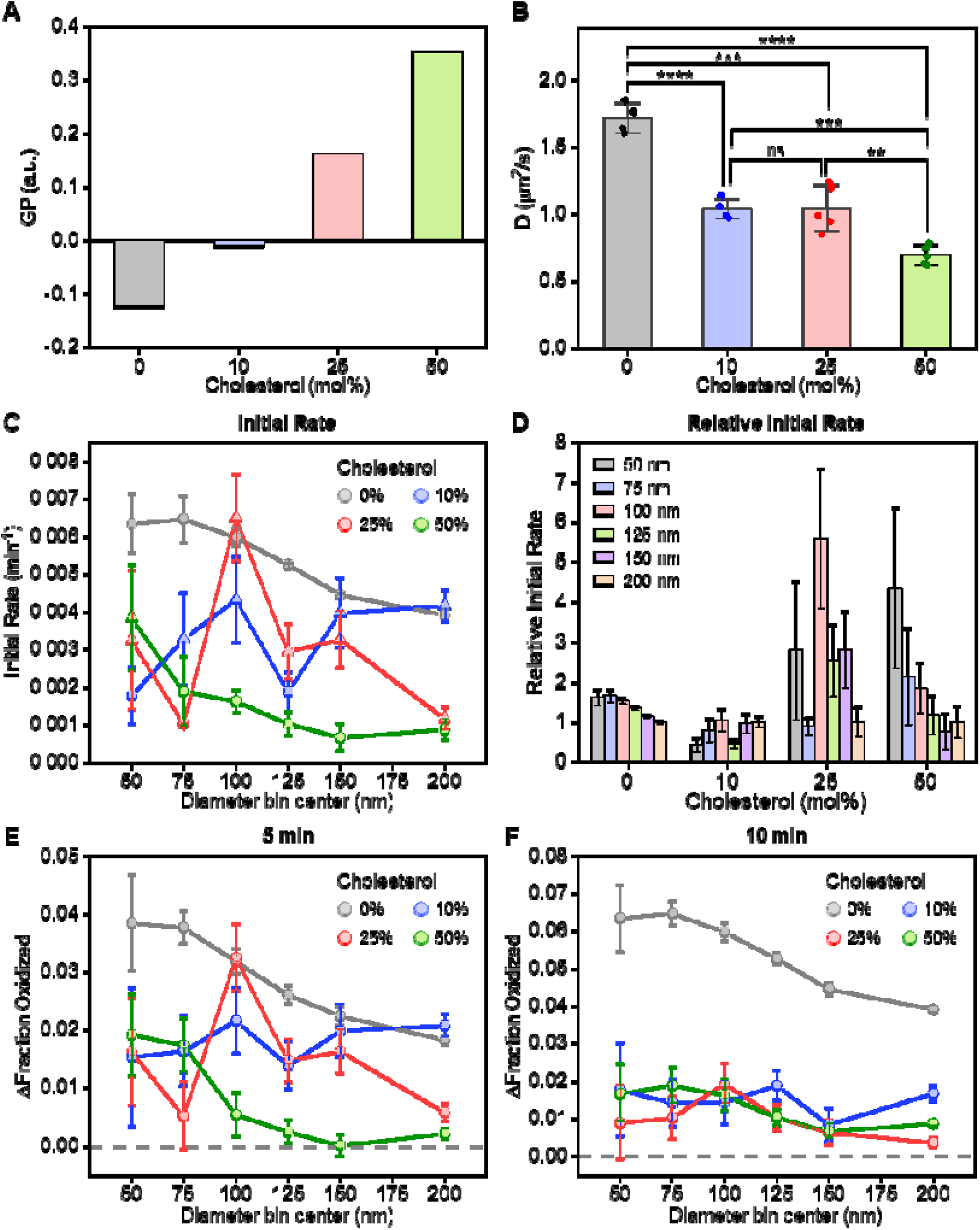
Cholesterol inhibits oxidation in DOPC membranes. **A)** The GP value of Laurdan for each membrane composition. **B)** Lateral diffusion coefficients for each membrane composition. **C)** The initial rate of oxidation as a function of vesicle diameter. **D)** The relative initial rate of oxidation in vesicles as a function of vesicle diameter for various concentration of cholesterol. The change in the fraction of oxidized C11-BODIPY after **E)** 5 minutes and **F)** 10 minutes of exposure to ROS, relative to its initial value for each diameter bin. Vesicles used in panel **A** were composed of DOPC (49.8-99.8 mol%), cholesterol (0-50 mol%), and Laurdan (0.2 mol%). SBLs used in panel **B** were composed of DOPC (49.875-99.875 mol%), cholesterol (0-50 mol%), and DPPE-ATTO 647N (0.125 mol%). Bars in **B** represent the mean and the error bars correspond to the standard deviation (*N*=4 for 0 mol% and 10 mol% cholesterol, and *N*=5 for 25 mol% and 50 mol% cholesterol, unpaired, two-sample t-test, ns: non-significance, **P<0.01, ***P<0.001, ****P<0.0001). All vesicles used in panel **C-F** were composed of DOPC (48.5-98.5 mol%), cholesterol (0-50 mol%), C11-BODIPY (0.5 mol%), DPPE-ATTO 647N (0.5 mol%), and DSPE-PEG(2000)-Biotin (0.5 mol%), and oxidation was induced using 7.5 μM H_2_O_2_ and 0.15 μM FeSO_4_. The number of SUVs in each diameter bin varied from 389–4003 (DOPC+10% Chol), 563–5124 (DOPC+25% Chol), and 585–5276 (DOPC+50% Chol). Data points in **C** represent the average initial rate of oxidation calculated from linear regression of three time points (0, 5, and 10 minutes, circle) or slope of the first two time points (0 and 5 minutes, triangle) and the error bars correspond to the standard error calculated from linear regression (circle) or propagated error from the standard error (triangle). Bars in **D** are calculated by normalizing initial rates for a given cholesterol mole percentage by the corresponding initial rate within 200 nm vesicles. Error bars represent the propagated uncertainty based on the errors of the underlying data sets. Data points in **E** and **F** represent the average value of the bin, and the error bars correspond to the standard error of difference. Each value displayed for a bin represents the midpoint of the bin. All bins were symmetric, with endpoints aligned to the start of the next bin, ensuring no overlap.

The inclusion of cholesterol reduced the initial oxidation rate in SUVs under most conditions (Figures 5C, S12A-D), although in some cases, the rates were comparable to cholesterol-free SUVs of the same diameter. Curvature sensitivity was evident in membranes lacking cholesterol but was diminished at 10 mol% and 25 mol% cholesterol (Figures 5D, S12E-H). At 50 mol% cholesterol, curvature sensitivity was restored and even exceeded that observed in cholesterol-free membranes. These trends were mirrored when examining changes in the fraction of oxidized C11-BODIPY after 5 minutes of ROS exposure (Figure 5E). For larger vesicles (i.e., > 100 nm diameter), the strongest suppression was observed at 50 mol% cholesterol. After 10 minutes of ROS exposure, oxidation was strongly suppressed in all cholesterol-containing SUVs (Figure 5F). At this later time point, the extent of oxidation suppression did not show a clear dependence on cholesterol content as all mole percentages exhibited similarly low levels of oxidation.

### Membrane curvature and lipid composition simultaneously impact oxidation in lipid bilayers

Membrane curvature, lipid packing, and lipid mobility are all factors that regulate oxidation in membranes (Figure 6A). Here we have consolidated the effects of membrane properties into a unified framework by examining oxidation rates as a function of membrane packing (GP), diffusivity (D), curvature, and lipid unsaturation (Figures 6B-E). Membrane curvature strongly modulates oxidation, with smaller vesicles (i.e., 50 nm diameter) exhibiting higher overall initial oxidation rates than larger vesicles (i.e., 200 nm diameter) (Figures 6B–C). Within each curvature regime, oxidation generally increases with increasing diffusivity and decreasing GP. This trend is consistent with increased lipid lateral mobility at higher diffusivity and enhanced lipid tail exposure and packing defects at lower GP, both of which facilitate oxidative reactions. This dependence is substantially more pronounced in highly curved membranes, where oxidation rates span a broader range across compositions, whereas in larger vesicles the variation is more compressed, indicating a weaker dependence on membrane packing and diffusivity.

**Figure 6.**
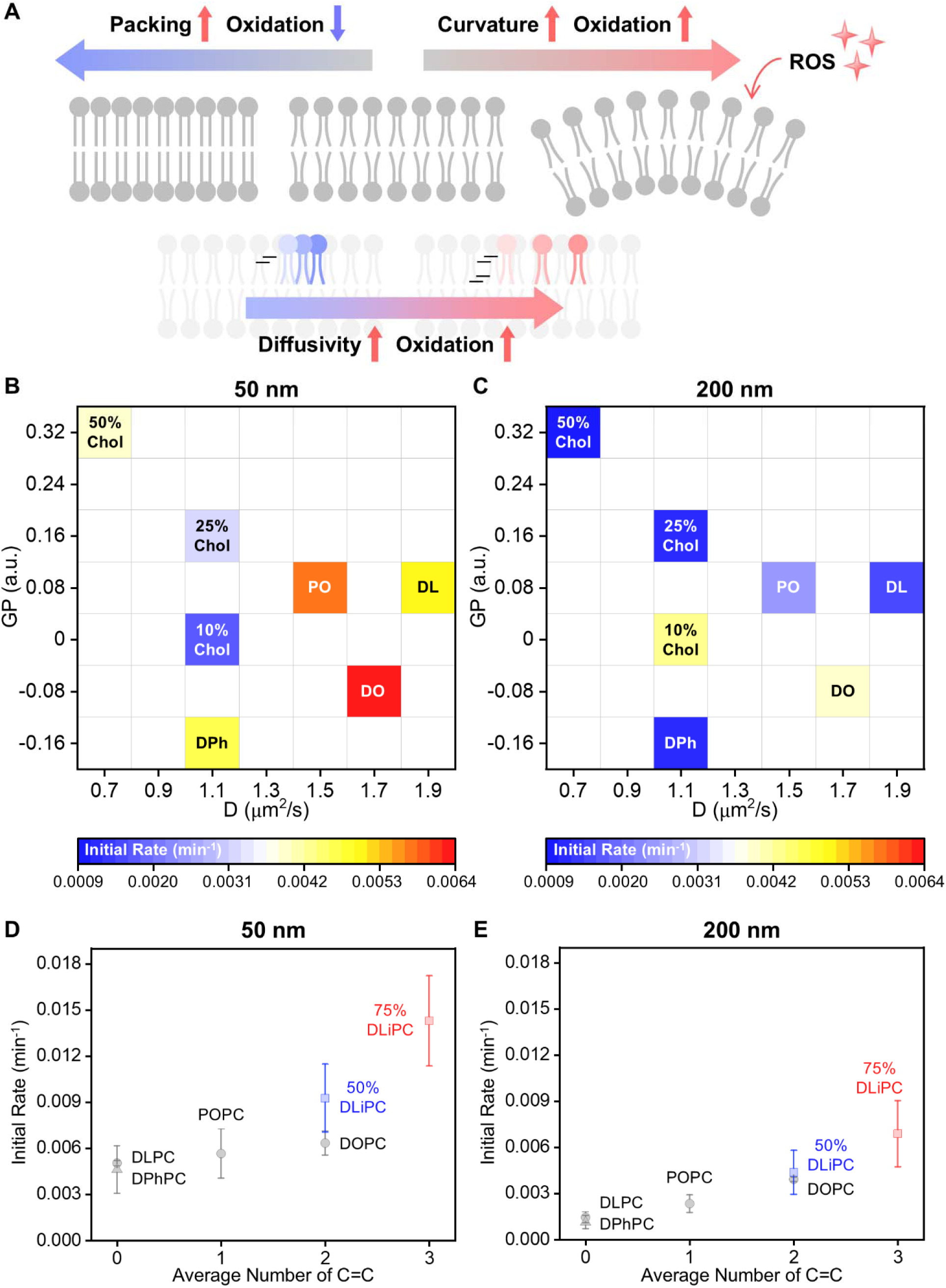
Impact of membrane curvature and lipid composition on oxidation in lipid bilayers. **A)** Schematic highlighting the impact of different membrane properties on oxidation in lipid bilayers. Heat map of initial oxidation rates as a function of membrane packing, GP, and diffusivity, D, in vesicle populations with average diameters of **B)** 50 nm and **C)** 200 nm. Initial oxidation rates as a function of the average number of carbon double bonds within lipids in vesicles with average diameters of **D)** 50 nm and **E)** 200 nm. Diffusion coefficients in panel B and C were obtained from SBLs. Lipid compositions are denoted as: DLPC (DL), DPhPC (DPh), POPC (PO), DOPC (DO), DOPC+10% Cholesterol (10% Chol), DOPC+25% Cholesterol (25% Chol), and DOPC+50% Cholesterol (50% Chol). Data points in panel D and E represent the average initial rate of oxidation and the error bars correspond to either the standard error calculated from linear regression or propagated error from the standard error.

The influence of lipid unsaturation further reinforces these trends. Increasing the number of carbon–carbon double bonds leads to higher oxidation rates in both curvature regimes (Figures 6D–E), with an approximately proportional increase in oxidation from one to three double bonds, approaching a threefold enhancement. This near-linear scaling is observed in both 50 nm and 200 nm vesicles, indicating that double bond content provides a consistent chemical driver of oxidation. However, the magnitude of this response is amplified in highly curved membranes, which exhibit higher absolute oxidation rates and a broader dynamic range compared to larger vesicles. Together, these results demonstrate that while unsaturation governs the baseline scaling of oxidation, membrane curvature enhances both the rate and sensitivity of this process.

## DISCUSSION

We developed a framework to systematically probe and quantify oxidation in lipid bilayers across a broad range of membrane environments. In nearly all membrane compositions examined, we observed a consistent trend in which C11-BODIPY underwent faster and more extensive oxidation in small, highly curved vesicles than larger ones. The shallow membrane position of C11-BODIPY (31, 38) likely accentuates this curvature-dependent response. This result suggests that membrane curvature amplifies the initiation step of oxidation, as highly curved membranes enhance lipid splay (i.e., increase lipid packing defects) (55), thereby facilitating greater flux of ROS into the hydrophobic region (21).

In addition to lipid packing defects, our results indicate that lipid mobility also plays a critical role in the oxidation process. Lipids in DPhPC membranes exhibited lower mobility than those in DLPC membranes (Figures 2I, S4), likely due to steric hindrance or tangling among the branched methyl chains (Figure 2A). Although DPhPC membranes are defect-rich, a property that enhances the initiation step of oxidation, they exhibited oxidative behavior that was comparable to that of DLPC membranes (Figures 2E-F, S9C). These findings suggest a compensatory relationship between the initiation and propagation steps of oxidation, in which membrane packing defects promote initiation while lipid mobility regulates propagation.

The propagation step can also be enhanced through the presence of carbon double bonds in unsaturated lipid tails. Even modest unsaturation, as in POPC with a single carbon double bond per lipid, significantly increased C11-BODIPY oxidation relative to DLPC and DPhPC membranes (Figures 3D-E, S9C), especially in membranes with low curvature (i.e., larger than 100 nm in diameter). While POPC and DOPC exhibited intermediate diffusivity and packing properties compared to DLPC and DPhPC (Figures 3B-C, S4, S6), the presence of double bonds provided additional chemical reactive sites that accelerated radical spreading through the membrane (10, 56). This finding indicates that chemical reactivity can outweigh physical membrane properties.

When comparing the 75% DLiPC lipid composition to DOPC, the rates and extents of oxidation in 75% DLiPC membranes were considerably higher for all vesicle sizes examined (Figures 4D-E), consistent with the higher susceptibility of PUFAs to oxidation (10) and the increased number of double bonds in the 75% DLiPC composition. The differences between DOPC and 50% DLiPC membranes, which both contain approximately two double bonds per phospholipid, were much less pronounced. This result is consistent 50% DLiPC membranes exhibiting both increased packing defects (i.e., lower GP) and reduced diffusivity (Figures 4B-C), where the pro-oxidative effect of greater tail exposure is counterbalanced by slower lipid mobility. Together, these results suggest that the density of double bonds sets the baseline susceptibility to oxidation, while membrane physical properties fine-tune the observed kinetics.

When considering cholesterol-free membranes, we found that the composition dependence of lipid oxidation was strongest in larger vesicles (> 100 nm diameter) but diminished at high curvature (∼ 50 nm diameter) (Figures 3D-E, 4D-F, 6B). At these small length scales, the abundance of packing defects promotes rapid initiation, which can dominate and mask compositional effects, whereas in larger vesicles, where defects are fewer and initiation is limited, lipid composition plays a more prominent role.

When present, cholesterol markedly suppressed oxidation (Figure 5), consistent with its ability to order lipid tails, decrease membrane permeability, and reduce lipid mobility (26, 57). Laurdan GP measurements confirmed that cholesterol monotonically reduced lipid tail exposure (i.e., packing defects) (Figures 5A, S11A), while FRAP showed a general decrease in diffusivity with increasing cholesterol content (Figures 5B, S11B). At high membrane curvature (< 100 nm diameter), all cholesterol-containing membranes inhibited oxidation relative to cholesterol-free membranes, although the response was not monotonic with cholesterol mole percentage (Figures 5C, E). At low curvature (> 100 nm diameter), 10 mol% and 25 mol% cholesterol produced similar levels of oxidation, whereas 50 mol% cholesterol consistently yielded the lowest levels of oxidation. In large vesicles with 50 mol% cholesterol, over the first 5 minutes of ROS exposure, the oxidation of C11-BODIPY was nearly completely suppressed, indicating a lag phase during the onset of oxidation (Figure 5E) (58). After 10 minutes of ROS exposure, oxidation levels were comparable for all mole percentages (Figure 5F). Interestingly, while 25 mol% cholesterol reduced packing defects compared to 10 mol% cholesterol, both compositions exhibited comparable diffusivity and similar oxidation outcomes. This result suggests that once defects are reduced beyond a threshold, further changes in curvature play a minimal role in initiation, and diffusivity becomes the dominant factor in oxidation. At 50 mol% cholesterol, diffusivity was further decreased, leading to a pronounced reduction in oxidation. Therefore, cholesterol reduces initiation by sealing packing defects that limit ROS access and suppresses propagation by decreasing lipid diffusivity.

While our results highlight the ability of cholesterol to reduce oxidation in DOPC membranes, this result may not be universal for all PC lipids. For example, dipalmitoylphosphocholine (DPPC) lipids, which form a tightly packed gel phase at room temperature, can become fluidized to a liquid-ordered phase upon the addition of cholesterol (26). In the context of our results, we would expect cholesterol to have an opposite effect in DPPC membranes where it enhances oxidation due to increases in hydrophobic tail exposure and membrane fluidity.

Cholesterol also altered the curvature sensitivity of oxidation (Figure 5D). In cholesterol-free membranes, oxidation rates increased with curvature, but this curvature dependence was lost at 10 mol% and 25 mol% cholesterol. At 50 mol% cholesterol, curvature sensitivity was restored and even exceeded the cholesterol-free condition, with a ∼4.3-fold difference between 50 nm and 200 nm vesicle populations compared to ∼1.6-fold without cholesterol. Given cholesterol’s strong influence on membrane permeability and lipid mobility, the observed changes in curvature sensitivity may result from its dual effects on oxidation. As cholesterol content increases, defect sealing could reach saturation rapidly, while reduced mobility continues to dampen propagation. However, at sufficiently high concentrations, cholesterol’s small -OH headgroup can create gaps between neighboring phospholipids that enlarge packing defect areas (59), effectively reintroducing defect-like features. Importantly, this effect occurs in both flat and curved membranes but is amplified in small, highly curved vesicles, where curvature already predisposes the bilayer to more pronounced defect formation. This curvature-dependent amplification provides a mechanistic explanation for how high cholesterol levels could reintroduce defect-like features and thereby restore curvature-sensitive oxidation.

## CONCLUSION

In conclusion, we developed a framework to examine how membrane curvature and lipid composition act together to regulate oxidation in lipid bilayers. Using C11-BODIPY and tethered vesicle assays, we showed that curvature enhances oxidation in lipid bilayers in a composition-dependent manner. When membranes are highly curved, the elevated density of packing defects promotes rapid initiation, accelerating oxidation and thereby diminishing compositional contrasts. On flatter membranes, where defect-mediated initiation is limited, oxidation becomes more sensitive to lipid chemistry, emphasizing the compositional control of oxidation. Beyond curvature, however, the extent of oxidation is governed by the interplay among several membrane properties, including defect density, lateral mobility, and chemical reactivity, which together determine how readily oxidation can initiate and propagate. Abundant defects alone do not guarantee greater oxidation if lateral mobility is restricted. The presence of carbon double bonds facilitates propagation and increases oxidation, with monounsaturated lipids enhancing oxidation and PUFA-containing lipids amplifying it further. Cholesterol suppresses oxidation by reducing tail exposure and mobility. At low to moderate cholesterol levels (10–25 mol%), the overall reductions in defect accessibility and lipid diffusivity diminish curvature sensitivity. At high cholesterol levels (50 mol%), diffusivity is slowed enough that curvature sensitivity re-emerges and even exceeds that of cholesterol-free membranes. Together, these findings demonstrate that oxidation in lipid bilayers reflects a balance between defect-driven initiation and mobility-driven propagation, tuned by both curvature and composition.

## Author contributions

J.K. and W.F.Z. designed experiments. J.K. and S.N.B. performed experiments. All authors consulted together on the interpretation of results and manuscript preparation.

## Data Availability

All data supporting our findings is available within the manuscript and supporting material. Raw data files can be shared upon reasonable request.

## Acknowledgements

^1^H NMR spectroscopy data was acquired from the Center of excellence for molecular characterization at the University of Southern California. This research was supported by the National Institutes of Health through R35GM147333 to J. Kim and W. F. Zeno.

