## Supporting Information for "Membrane curvature enhances oxidation within lipid bilayers in a composition-dependent manner"

### Supporting Discussion

#### 1. Conversion of C11-BODIPY fluorescence intensity into concentration

The fluorescence emission spectra of C11-BODIPY in its initial and fully oxidized states were acquired by spectrofluorometer (Figure S2). First, the fully oxidized C11-BODIPY spectrum was integrated over the range of 500–540 nm (E520), which is the same detection range used in fluorescence microscopy (Figure S2B, blue region). As the concentration of oxidized C11-BODIPY ($C_{ox}$) is equal to the total concentration of C11-BODIPY ($C_{total}$), the fluorescence intensity coefficient of oxidized C11-BODIPY (ε520) was calculated using equation S1. Next, from the initial C11-BODIPY spectrum, the concentration of oxidized C11-BODIPY was calculated (Figure 2SA, blue region). Using equation S2, the concentration of unoxidized C11-BODIPY was calculated ($C_{unox}$). By integrating over the range of 570–620 nm (E590), and using equation S3, ε590 was calculated (Figure S2A, gray patterned region). By using the obtained fluorescence intensity coefficients and substituting E520 and E590 with the fluorescence intensities measured from the tethered vesicle assay, the molar fraction of oxidized C11-BODIPY was calculated (equation S4).

E520: integrated area of C11-BODIPY emission spectrum from 500 to 540 nm

E590: integrated area of C11-BODIPY emission spectrum from 570 to 620 nm

ε520: fluorescence intensity coefficient of oxidized C11-BODIPY

ε590: fluorescence intensity coefficient of unoxidized C11-BODIPY

$C_{ox}$: concentration of oxidized C11-BODIPY

$C_{unox}$: concentration of unoxidized C11-BODIPY

$$E520 = \varepsilon 520 \times C_{ox} \qquad \text{eq (S1)}$$

$$C_{total} = C_{ox} + C_{unox} \qquad \text{eq (S2)}$$

$$E590 = \varepsilon 590 \times C_{unox} \qquad \text{eq (S3)}$$

$$\frac{C_{ox}}{C_{total}} = \frac{E520/\varepsilon 520}{E520/\varepsilon 520 + E590/\varepsilon 590} \qquad \text{eq (S4)}$$

### 2. Calculating diffusion coefficients from FRAP

To control for additional photobleaching during fluorescence recovery, a bilayer region in close proximity to the bleached region was used to normalize fluorescence intensities. This neighboring region was of identical size to the bleached region (15 μm in diameter) and not subjected to 100% laser intensity. The fluorescence intensity of this control, neighboring region, $F_{max}(t)$, was monitored over time. The mean fluorescence intensity of the bleached region, F(t), was normalized using equation S5. An isotropic lateral diffusion system can be described by the diffusion equation (equation S6), and using a uniform circular beam model, the normalized intensity was fitted using equation S7 (1, 2). Finally, the diffusion coefficient, D, was calculated using equation S8.

$F_N(t)$: normalized intensity at time=t

$F(t)$: mean intensity of the bleached region at time=t

$F_0$: mean intensity of the bleached region immediately after bleaching

$F_{max}(t)$: mean background intensity at time=t

A: fitting parameter

$\tau_D$: characteristic diffusion time

$I_0, I_1$: modified Bessel function of $0^{th}$ and $1^{st}$ order

$w$: radius of the bleached circle

$$F_N(t) = \frac{F(t) - F_0}{F_{max}(t) - F_0} \qquad \text{eq (S5)}$$

$$\frac{\partial C(r,t)}{\partial t} = D\left\{\frac{1}{r}\frac{\partial C(r,t)}{\partial r} + \frac{\partial^2 C(r,t)}{\partial r^2}\right\} \qquad \text{eq (S6)}$$

Boundary conditions: symmetry at $r = 0$, $C(w,t) = C_0$ ($C_0$: initial concentration of the fluorophore)

Initial condition: $C(r,0) = 0$

$$F_N(t) = A \cdot \exp(-\frac{2\tau_D}{t}) \cdot \left\{I_0\left(\frac{2\tau_D}{t}\right) + I_1\left(\frac{2\tau_D}{t}\right)\right\} \qquad \text{eq (S7)}$$

$$D = w^2/4\tau_D \qquad \text{eq (S8)}$$

**3. Determining the effective lipid concentration used in the tethered vesicle assay**

To determine the effective concentration of small unilamellar vesicles (SUVs) used in our tethered vesicle assays, the concentration of the tethered SUVs on a surface was converted into the concentration of free SUVs in solution. First, the effective volume of individual vesicle ($V_{eff}$) was calculated using equation S9. The area of image frame refers to the size of a single frame taken by confocal microscope (71.52×71.52 $\mu m^2$) and the average number of vesicles per frame was calculated using CME analysis (3). A total of 10 frames were taken per experimental condition. The effective concentration ($C_{eff}$) was calculated using equation S10, where r is the average radius of SUVs, lipid area refers to the projected area on the membrane (0.6 $nm^2$) (4) and $N_A$ is Avogadro's number. The converted effective concentration of SUVs was used to calculate the concentration of Fenton's reagent required for oxidation in free volume.

$$V_{eff} = \left(\frac{\text{Area of image frame}}{\text{Average number of vesicles per frame}}\right)^{1.5} \quad \text{eq (S9)}$$

$$C_{eff} = \frac{2\times(4\pi r^2)}{\text{Lipid area}\times V_{eff}\times N_A} \quad \text{eq (S10)}$$

| Lipid composition | Z-average (nm) |
|---|---|
| DLPC (TVA) | 83.8 ± 1.2 |
| DPhPC (TVA) | 137.4 ± 1.1 |
| POPC (TVA) | 102.0 ± 2.5 |
| DOPC (TVA) | 112.0 ± 0.8 |
| DLiPC:DPhPC (1:1) (TVA) | 112.0 ± 1.6 |
| DLiPC:DPhPC (3:1) (TVA) | 106.9 ± 1.6 |
| DOPC+Cholesterol 10 mol% (TVA) | 112.1 ± 0.8 |
| DOPC+Cholesterol 25 mol% (TVA) | 122.2 ± 1.7 |
| DOPC+Cholesterol 50 mol% (TVA) | 132.1 ± 3.2 |
| POPC (NMR) | 132.3 ± 1.9 |
| DOPC (NMR) | 120.8 ± 1.6 |
| DLPC:Laurdan (99.8:0.2) | 160.5 ± 2.1 |
| DPhPC:Laurdan (99.8:0.2) | 141.7 ± 0.8 |
| POPC:Laurdan (99.8:0.2) | 162.8 ± 2.8 |
| DOPC:Laurdan (99.8:0.2) | 147.1 ± 1.9 |
| DLiPC:DPhPC:Laurdan (49.9:49.9:0.2) | 152.2 ± 1.7 |
| DLiPC:DPhPC:Laurdan (74.85:24.95:0.2) | 150.7 ± 0.8 |
| DOPC:Cholesterol:Laurdan (89.8:10:0.2) | 148.2 ± 2.8 |
| DOPC:Cholesterol:Laurdan (74.8:25:0.2) | 159.8 ± 2.3 |
| DOPC:Cholesterol:Laurdan (49.8:50:0.2) | 141.2 ± 2.9 |
| DLPC (Anisotropy) | 133.7 ± 3.2 |
| DPhPC (Anisotropy) | 129.4 ± 1.7 |
| POPC (Anisotropy) | 124.9 ± 0.9 |
| DOPC (Anisotropy) | 118.0 ± 0.8 |

**Table S1. DLS measurements of SUVs of each lipid composition.** Vesicles used for tethered vesicle assays (TVA) are composed primarily of PC lipids listed in the table and trace amounts of C11-BODIPY, DPPE-ATTO 647N, and DSPE-PEG(2000)-Biotin (*N*=3, average ± standard deviation).

| Lipid composition | $\varepsilon_{520}/\varepsilon_{590}$ |
|---|---|
| DLPC | 5.26 |
| DPhPC | 4.38 |
| POPC | 3.94 |
| DOPC | 4.64 |
| DLiPC:DPhPC (1:1) | 3.72 |
| DLiPC:DPhPC (3:1) | 2.89 |
| DOPC+Cholesterol 10 mol% | 3.96 |
| DOPC+Cholesterol 25 mol% | 4.17 |
| DOPC+Cholesterol 50 mol% | 4.38 |

**Table S2. Fluorescence intensity coefficient ratios for each lipid composition.**

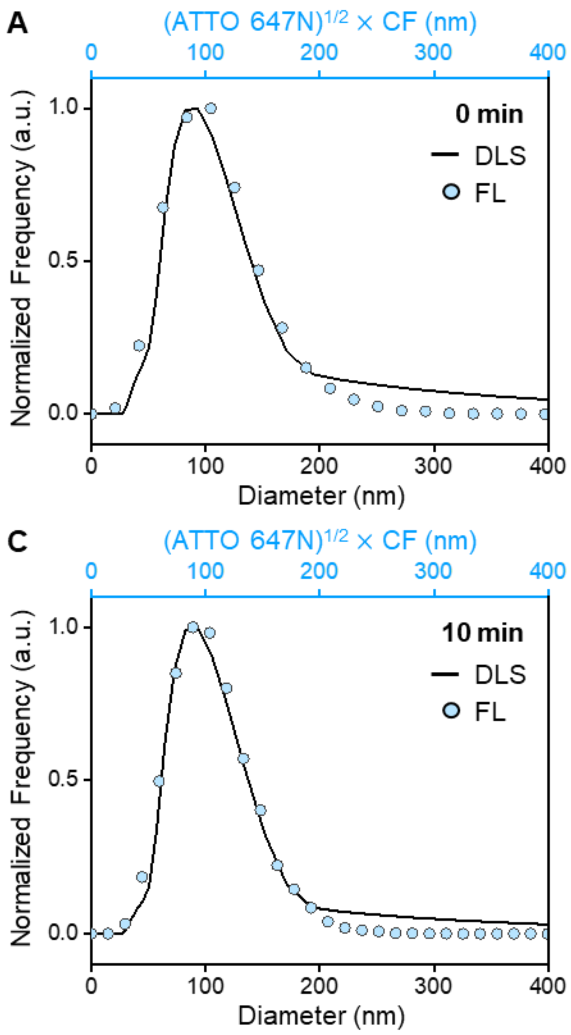


**Figure S1. Conversion of fluorescence intensity to vesicle diameter.** Calibration of the ATTO 647N fluorescence intensity with the diameter of vesicles that were incubated with Fenton's reagent for **A)** 0, **B)** 5, **C)** 10, and **D)** 30 minutes. Vesicles were composed of DLPC (98.5 mol%), C11-BODIPY (0.5 mol%), DPPE-ATTO 647N (0.5 mol%), and DSPE-PEG(2000)-Biotin (0.5 mol%), and oxidation was induced using 7.5 µM $H_2O_2$ and 0.15 µM $FeSO_4$. The size distribution was obtained by dynamic light scattering (DLS), and the fluorescence intensity of ATTO 647N was obtained by fluorescence microscopy based tethered vesicle assay. The conversion factor (CF) was calculated by overlaying the size distribution with the square root of the fluorescence intensity (black line: DLS measurement, blue dot: square root of the fluorescence intensity of ATTO 647N multiplied by CF).

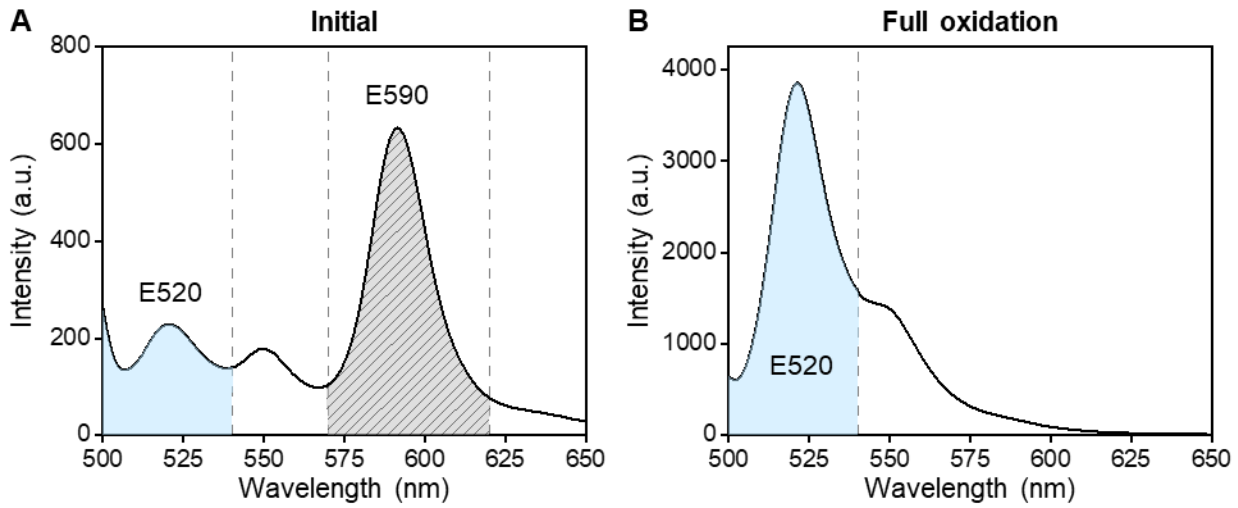


**Figure S2. Emission spectra of C11-BODIPY in its initial and fully oxidized states.** Representative fluorescence emission spectra of C11-BODIPY in **A)** its initial state and **B)** its fully oxidized state. Vesicles were composed of 99.95 mol% DLPC and 0.05 mol% C11-BODIPY (200 μM total lipid concentration), and oxidation was induced using 20 mM $H_2O_2$ and 1 mM $FeSO_4$. Each spectrum was integrated over the range of 500–540 nm to obtain oxidized C11-BODIPY intensity (blue region, E520), and 570–620 nm for unoxidized C11-BODIPY intensity (gray patterned region, E590).

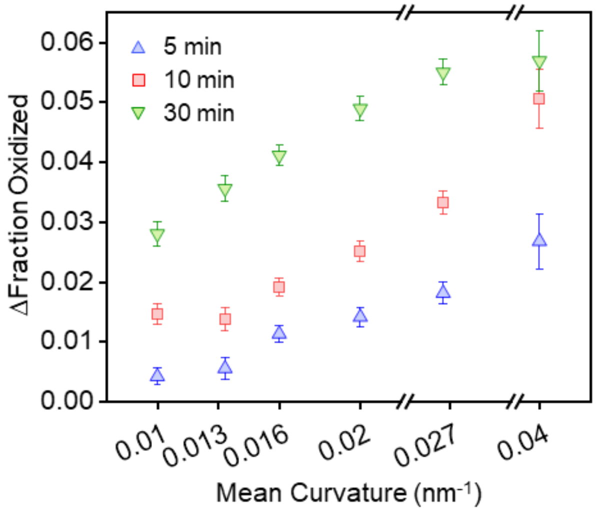


**Figure S3. Greater fractions of C11-BODIPY get oxidized as membrane curvature increases.** The change in the molar fraction of oxidized C11-BODIPY relative to its initial value for each membrane curvature bin for different experimental time points. The mean curvature was calculated by the inverse of the vesicle radius. Vesicles were composed of DLPC (98.5 mol%), C11-BODIPY (0.5 mol%), DPPE-ATTO 647N (0.5 mol%), and DSPE-PEG(2000)-Biotin (0.5 mol%), and oxidation was induced using 7.5 μM $H_2O_2$ and 0.15 μM $FeSO_4$. Data points represent the average value of the bin, and the error bars correspond to the standard error of the mean for each bin. Each value displayed for a bin represents the midpoint of the bin. All bins were symmetric, with endpoints aligned to the start of the next bin, ensuring no overlap.

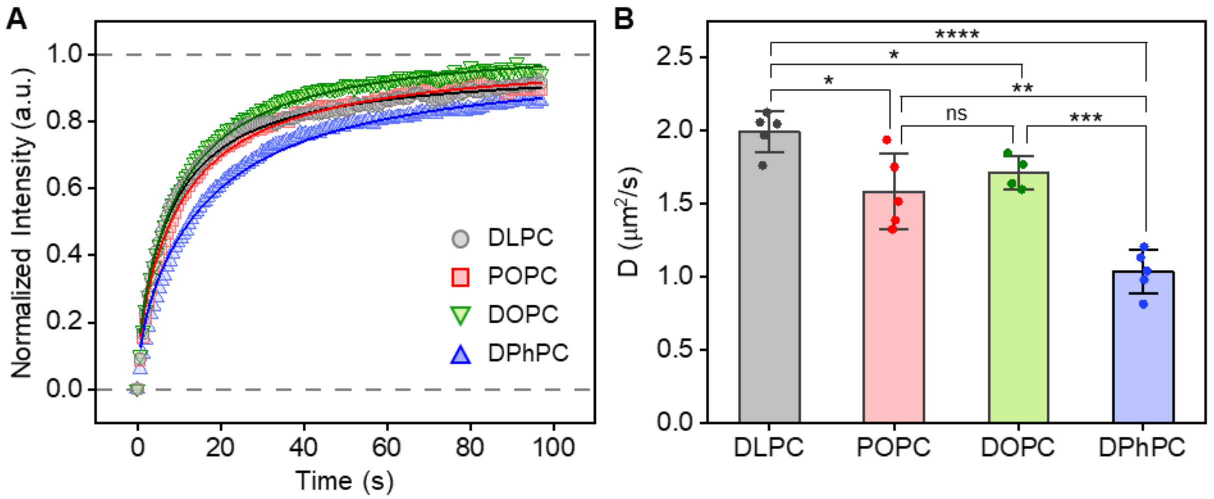


**Figure S4. FRAP on SBLs composed of various lipid compositions. A)** Recovery of fluorescence in SBLs over time. The lines represent the best fits to equation S7. **B)** Diffusion coefficients for each lipid composition. SBLs were composed of PC lipids (99.875 mol%) and DPPE-ATTO 647N (0.125 mol%). Data points in **A** represent the average normalized fluorescence intensity. Bars in **B** represent the mean and the error bars correspond to the standard deviation (*N*=5 for DLPC, POPC, and DPhPC, *N*=4 for DOPC, unpaired, two-sample t-test, ns: non-significance, *$P<0.05$, **$P<0.01$, ***$P<0.001$, ****$P<0.0001$).

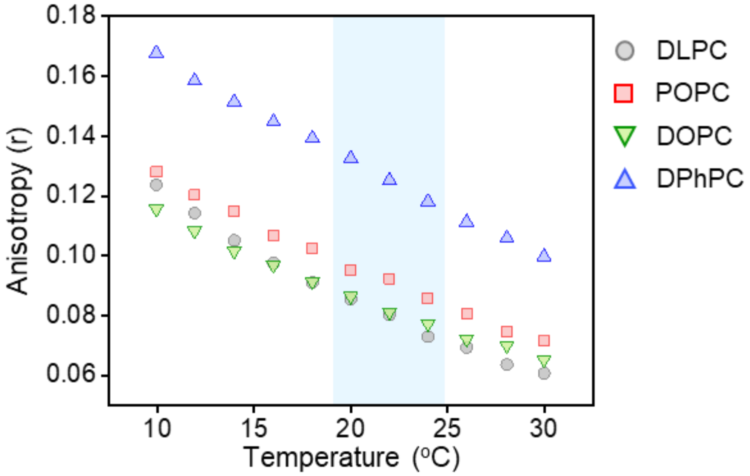


**Figure S5. Fluorescence anisotropy of DPH with various lipid compositions.** The fluorescence anisotropy of DPH in vesicles as a function of temperature. Vesicles were composed of PC lipids (60 μM total lipid concentration), and DPH was added at a molar ratio of 100:1 lipid:DPH. All oxidation experiments were conducted at room temperature (region highlighted in blue).

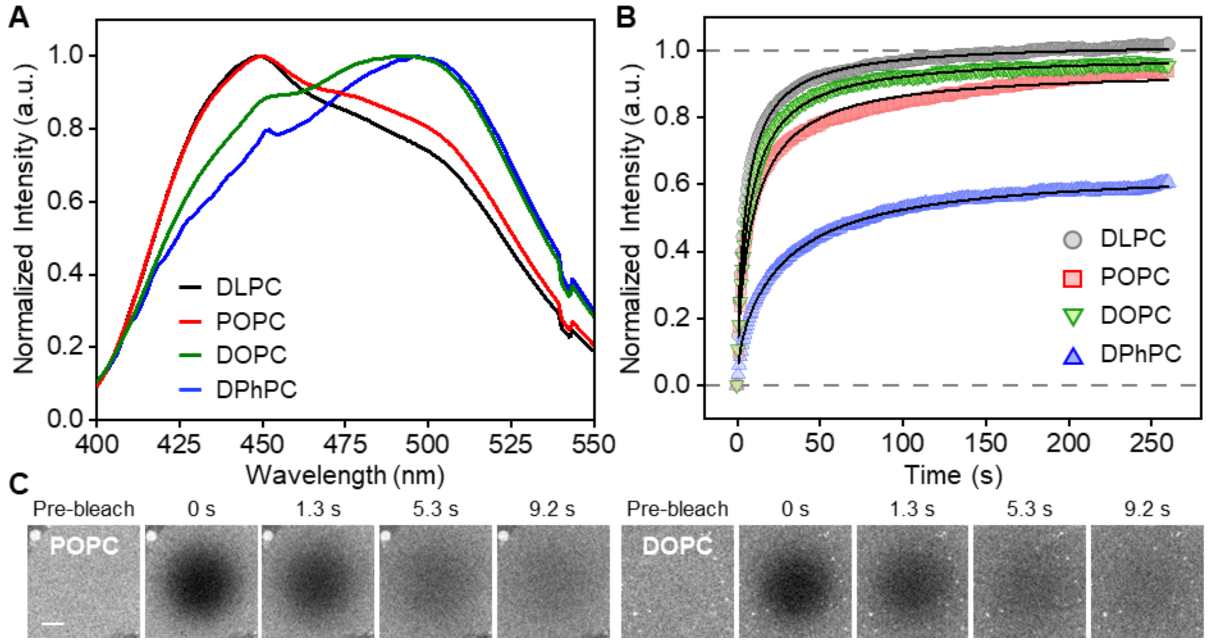


**Figure S6. Physical properties of the membranes composed of various lipid compositions. A)** The emission spectra of Laurdan within each lipid composition. **B)** Fluorescence recovery in MBLs over time. The black lines represent the lines of best fit from equation S7. **C)** Representative fluorescence micrographs of MBLs before and after photobleaching (scale bar: 5 μm). Vesicles used in **A** were composed of PC lipids (99.8 mol%) and Laurdan (0.2 mol%). MBLs used in **B-C** were composed of PC lipids (99.875 mol%) and DPPE-ATTO 647N (0.125 mol%). Data points in **B** represent the average normalized fluorescence intensity.

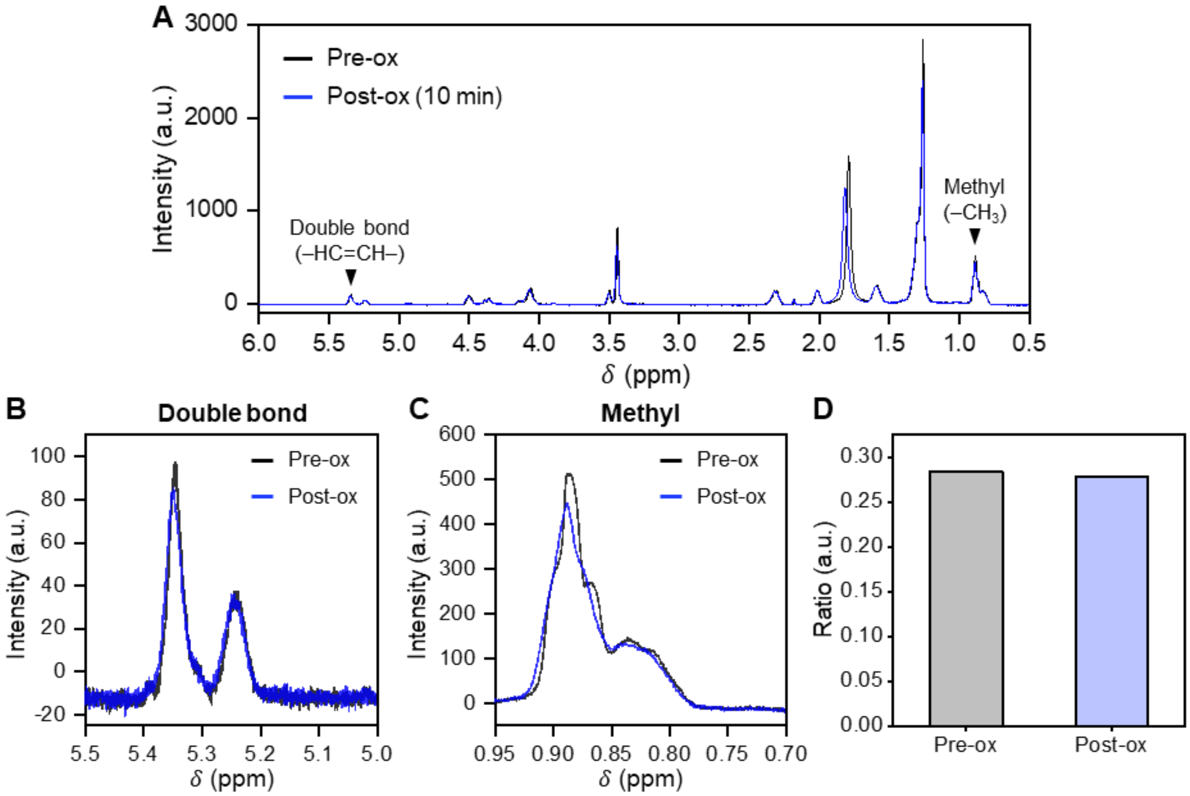


**Figure S7. [1]H NMR spectra of POPC before and after oxidation. A)** The full [1]H NMR spectra of POPC before oxidation (Pre-ox) and after 10 minutes of oxidation (Post-ox). The enlarged plots of **B)** the double bond and **C)** the methyl group peak of each condition. **D)** The ratio of the area under the double bond peak to the methyl group peak. Vesicles were composed of 100% POPC (2 mM total lipid concentration), and oxidation was induced using 320 μM $H_2O_2$ and 6.4 μM $FeSO_4$.

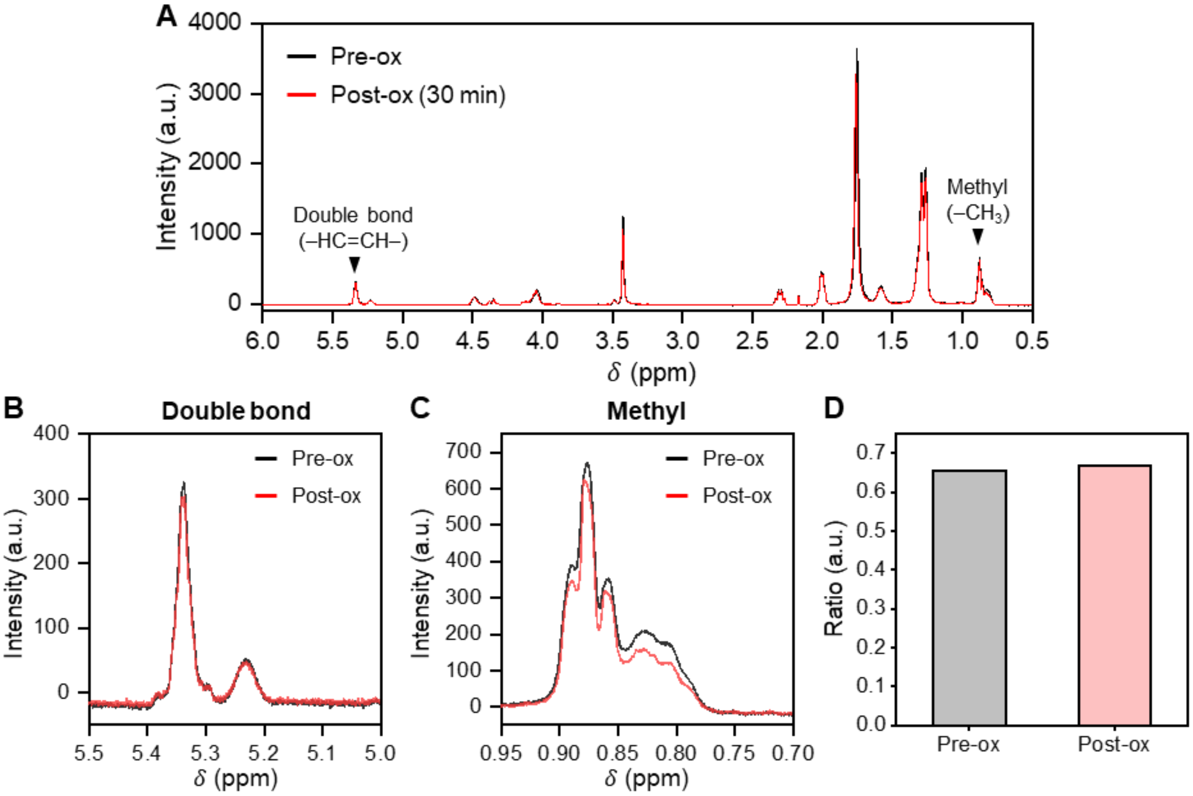


**Figure S8. [1]H NMR spectra of DOPC before and after oxidation. A)** The full [1]H NMR spectra of DOPC before oxidation (Pre-ox) and after 30 minutes of oxidation (Post-ox). The enlarged plots of **B)** the double bond and **C)** the methyl group peak of each condition. **D)** The ratio of the area under the double bond peak to the methyl group peak. Vesicles were composed of 100% DOPC (2 mM total lipid concentration), and oxidation was induced using 309 µM $H_2O_2$ and 6.2 µM $FeSO_4$.

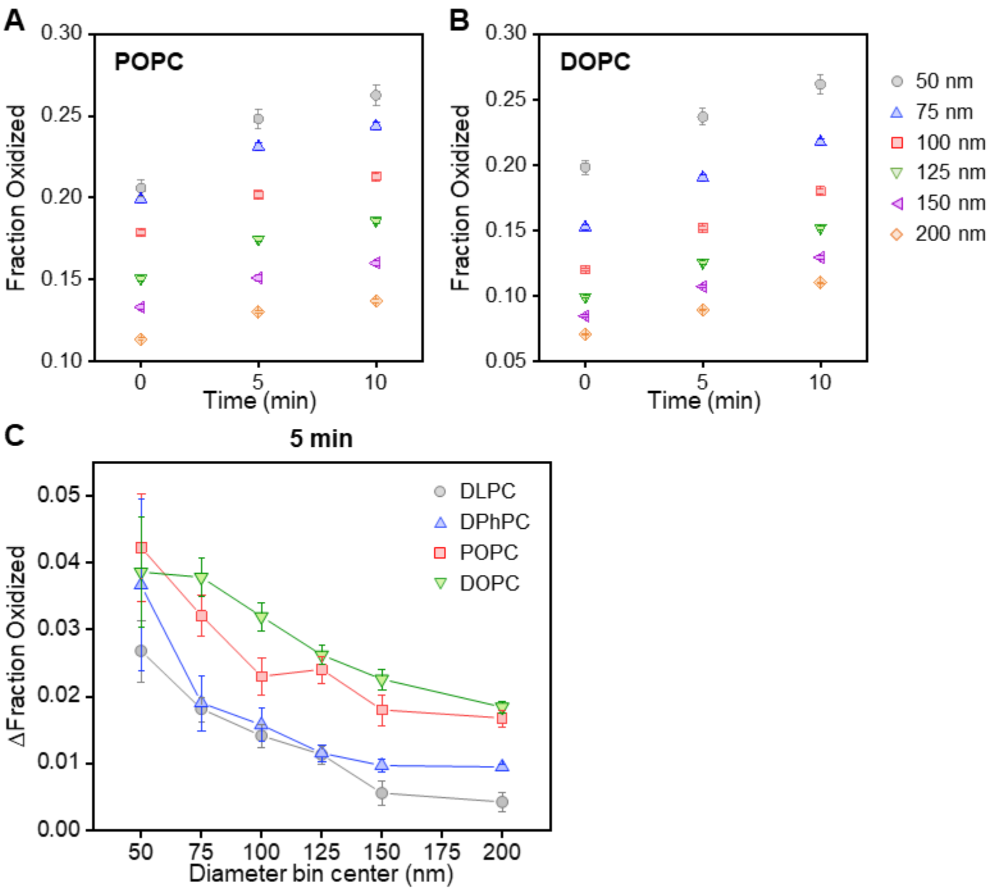


**Figure S9. Oxidation of C11-BODIPY in the presence of monounsaturated lipids.** The average fraction of oxidized C11-BODIPY in **A)** POPC and **B)** DOPC vesicles in each diameter bin over time. **C)** The change in the fraction of oxidized C11-BODIPY after 5 minutes of oxidation, relative to its initial value for each diameter bin. Vesicles were composed of POPC or DOPC (98.5 mol%), C11-BODIPY (0.5 mol%), DPPE-ATTO 647N (0.5 mol%), and DSPE-PEG(2000)-Biotin (0.5 mol%), and oxidation was induced using 7.5 µM $H_2O_2$ and 0.15 µM $FeSO_4$. Data points represent the average value of the bin, and the error bars correspond to the standard error of the mean for each bin. Each value displayed for a bin represents the midpoint of the bin. All bins were symmetric, with endpoints aligned to the start of the next bin, ensuring no overlap.

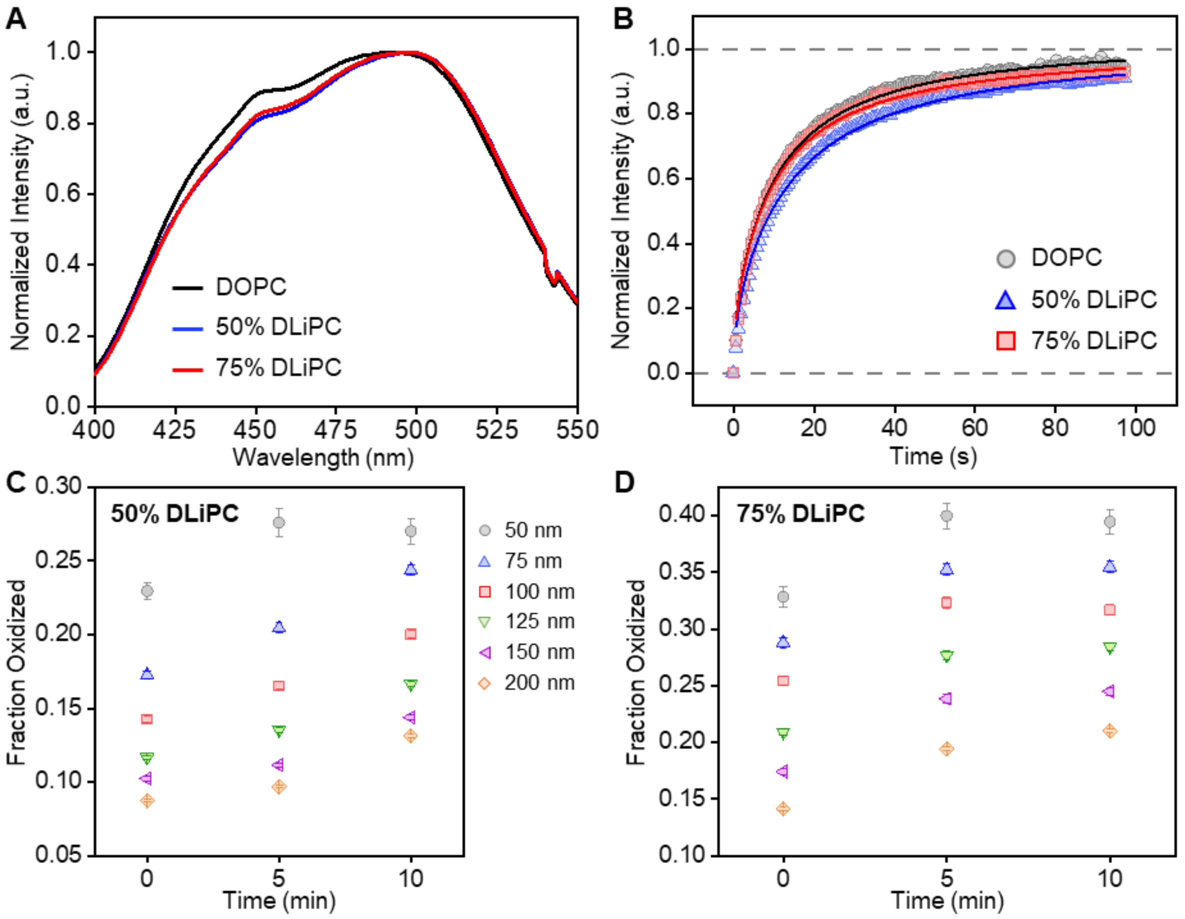


**Figure S10. Physical properties of the membranes and oxidation of C11-BODIPY in the presence of polyunsaturated lipids. A)** The emission spectra of Laurdan within each lipid composition. **B)** Recovery of fluorescence in SBLs over time. The lines represent the best fits with equation S7. The average fraction of oxidized C11-BODIPY in **C)** 50% DLiPC and **D)** 75% DLiPC vesicles in each diameter bin over time. Vesicles used in **A** were primarily composed of DOPC, 1:1 DLiPC:DPhPC (50% DLiPC), or 3:1 DLiPC:DPhPC (75% DLiPC), and Laurdan (0.2 mol%). SBLs used in **B** were primarily composed of DOPC, 1:1 DLiPC:DPhPC (50% DLiPC), or 3:1 DLiPC:DPhPC (75% DLiPC), and contained 0.125 mol% DPPE-ATTO 647N. Data points in **B** represent the average normalized fluorescence intensity. Vesicles used in **C** and **D** were primarily composed of 1:1 DLiPC:DPhPC (50% DLiPC) or 3:1 DLiPC:DPhPC (75% DLiPC) and contained C11-BODIPY (0.5 mol%), DPPE-ATTO 647N (0.5 mol%), and DSPE-PEG(2000)-Biotin (0.5 mol%). Oxidation was induced using 7.5 µM $H_2O_2$ and 0.15 µM $FeSO_4$. Data points in **C** and **D** represent the average value of the bin, and the error bars correspond to the standard error of the mean for each bin. Each value displayed for a bin represents the midpoint of the bin. All bins were symmetric, with endpoints aligned to the start of the next bin, ensuring no overlap.

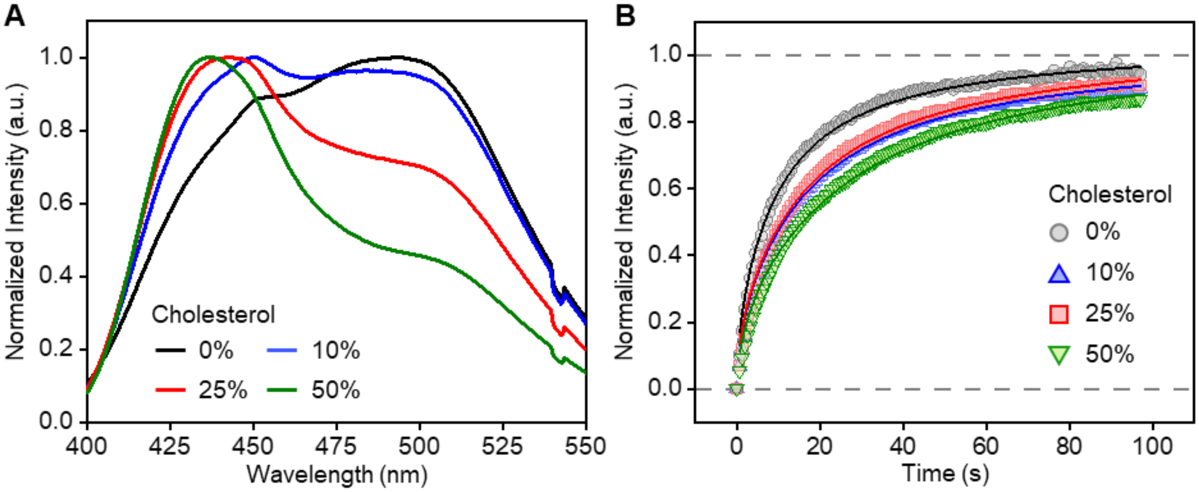


**Figure S11. Physical properties of the membranes composed of DOPC and cholesterol. A)** The emission spectra of Laurdan for each membrane composition. **B)** Fluorescence recovery in SBLs over time. The lines represent the best fits to equation S7. Vesicles used in **A** were composed of DOPC (49.8-99.8 mol%), cholesterol (0-50 mol%), and Laurdan (0.2 mol%). SBLs used in **B** were composed of DOPC (49.875-99.875 mol%), cholesterol (0-50 mol%), and DPPE-ATTO 647N (0.125 mol%). Data points in **B** represent the average normalized fluorescence intensity.

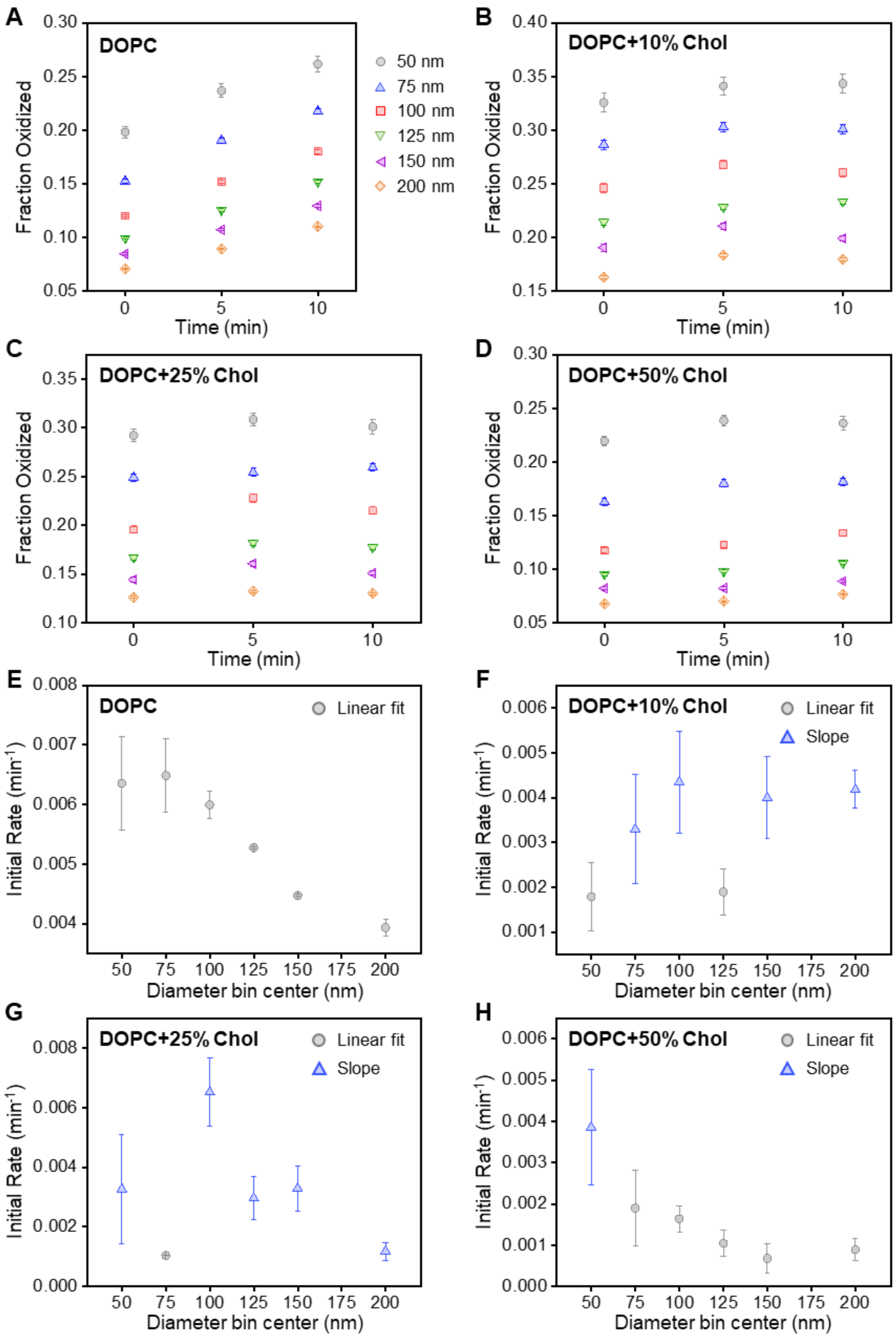


**Figure S12. Oxidation of C11-BODIPY in membranes composed of DOPC and various cholesterol**

**concentrations.** The average fraction of oxidized C11-BODIPY in the presence of **A)** 0 mol%, **B)** 10 mol%, **C)** 25 mol%, and **D)** 50 mol% cholesterol in each diameter bin over time. The initial rate of oxidation in the presence of **E)** 0 mol%, **F)** 10 mol%, **G)** 25 mol%, and **H)** 50 mol% cholesterol as a function of vesicle diameter. Vesicles were composed of DOPC (48.5–98.5 mol%), cholesterol (0–50 mol%), C11-BODIPY (0.5 mol%), DPPE-ATTO 647N (0.5 mol%), and DSPE-PEG(2000)-Biotin (0.5 mol%), and oxidation was induced using 7.5 μM $H_2O_2$ and 0.15 μM $FeSO_4$. Data points in **A-D** represent the average value of the bin, and the error bars correspond to the standard error of the mean for each bin. Data points in **E-H** represent the average initial rate of oxidation calculated from linear regression of three time points (0, 5, and 10 minutes, grey circle) or slope of the first two time points (0 and 5 minutes, blue triangle). The error bars correspond to the standard error calculated from linear regression (grey) or propagated error from the standard error (blue). Each value displayed for a bin represents the midpoint of the bin. All bins were symmetric, with endpoints aligned to the start of the next bin, ensuring no overlap.
